# Metabolic reprogramming of methylthioadenosine-dependent sulfur recycling is a major driver of CHIKV infection

**DOI:** 10.1101/2025.07.11.664323

**Authors:** Jackson M. Muema, Myriam Friemelt, Heike Overwin-Moser, Ulrike Beutling, Raimo Franke, Mark Brönstrup, Peter Sandner, Ursula Bilitewski

## Abstract

The supply of key metabolites into viral replication compartments must be assured through a coordinated reprogramming of host metabolic pathways. For chikungunya virus (CHIKV), the cellular metabolites required for a successful infection are largely unknown. We show that CHIKV reprograms sulfur-dependent pathways. To maintain the resupply of thiols, the methionine (Met) salvage players, 5ʹ-methylthioadenosine (MTA) and methionine adenosyltransferase-2a (Mat2a) are co-induced specifically. Under sulfur-depleted conditions, exogenously added MTA restores CHIKV replication more efficiently than its precursor *S*-adenosylmethionine, while inhibitions of Mat2a or *de novo* cysteine (Cys) biosynthesis reduce viral infectivity. Upon sulfur insufficiency CHIKV upregulates the U_34_-tRNA methyltransferase ALKBH8, and when ALKBH8 is deleted, virus replication is reduced by impairing sulfur relay, recapitulating Met-Cys deprivation effects. We found that MTA-mediated CHIKV replication occurs via m^6^A-independent priming, and that the *S*-adenosylhomocysteine hydrolase inhibitors DzNep and Adox, inhibited CHIKV replication with nanomolar potency. Our findings uncovered a major pro-CHIKV metabolic rheostat regulating tRNA modifications that can be targeted with host-directed antiviral agents.

## Introduction

Chikungunya virus (CHIKV) is a positive-strand RNA alphavirus (family: *Togaviridae*) first reported ∼75 years ago in the Newala district of Tanzania^1^. Since then, the virus, which is transmitted via an infectious bite of a female *Aedes* spp mosquito, has emerged and re-emerged intermittently in more than 100 countries across Africa, the Americas, Europe, Middle East and Pacific Island regions, usually causing mild febrile illnesses but also chronically debilitating arthralgic musculoskeletal syndrome. Fatalities from CHIKV infection are fortunately rare, accounting for 0.1% cases in high-risk areas, and are associated with multi-organ failure and dysregulated immune responses^2^. The lack of effective CHIKV treatments, coupled to the adaptations of *Aedes* mosquito vector allowing expansion to new ecologies arising from the dreadful impacts of climate change, raises increasing concerns^3–6^. As a consequence, over three quarters of the total world population is at-risk of CHIKV infections, with a strong warning of possible future pandemic outbreaks already predicted by the newly formed WHO Arbovirus Initiative^7^.

A major hallmark of viral infection encompasses hijacking of key cellular pathways to mobilize crucial resources into genome replication compartments^8–10^. Rapidly replicating RNA viruses infecting human cells, for instance SARS-CoV-2, Zika virus (ZIKV), and Dengue virus (DENV), have been described to actively reprogram specific metabolic pathways of host cells for a successful life cycle^11–16^ and pathogenesis^17–21^. However, despite several years of intensive investigations on CHIKV, it has remained unclear how the virus specifically reprograms host cell metabolism and most of the infection aspects are still derived from other closely related alphaviruses.

Because amino acids metabolism plays a vital role in providing the major biosynthetic precursors of proteins, nucleotides, and lipids, as well as their specialized regulatory intermediates, it is often hijacked during human viral infections^22–26^. As such, specific amino acids including glycine, threonine, serine, arginine, tryptophan, aspartate, glutamine, and asparagine have already been described to promote viral replications of SARS-CoV-2, and Influenza virus, or to serve as serum diagnostic viral biomarkers^27^. In eukaryotic cells, the sulfur-containing amino acids methionine (Met) and cysteine (Cys) provide essential thiol substrates of the Met arm of the one-carbon (1C) metabolism cycle that drive RNA processing^28,29^ and of iron-sulfur (Fe-S) clusters biogenesis^30^. Specifically, the synthesis of *S*-adenosylmethionine (SAM), the major methyl donor for posttranscriptionally modified RNAs, from Met catalyzed by the methionine adenosyltransferase 2a (Mat2a) is tightly dependent on Met availability, and its recovery is metabolically costly. Mat2a senses cellular Met/SAM levels and transcriptionally coordinates its regulatory function with 5ʹ-methylthioadenosine (MTA), a hydrophobic sulfur intermediate metabolite from the Met salvage cycle described to intersect with signaling pathways for cellular functions. Though not previously described for CHIKV, an interaction of Mat2a with viral RNA has been reported being pro-viral^31–33^. The contribution of these sulfur-containing amino acids to viral RNA replications has for long remained undefined.

Recently, studies of CHIKV-specific mechanisms facilitating vital RNA processing and translation steps have started^34^. To-date only two independent studies^35,36^ have demonstrated CHIKV to depend on host cell lipid metabolism, whereas knowledge on the functional relevance of other major metabolic pathways is largely missing. To address this critical knowledge gap, we studied CHIKV-infected cells applying untargeted metabolomics. We specifically detected an increased 5ʹ-methylthioadenosine (MTA) level, which was accompanied by increased methionine adenosyltransferase 2a (Mat2a). Our data further show that, in the absence of sulfur-containing amino acids, exogenously-added MTA is a direct-acting pro-CHIKV metabolite that also exerts its effects at transcriptional level of viral RNA replication. Further exploration of CHIKV dependency on sulfur indicates that this virally-induced metabolic reprogramming is therapeutically targetable and links tRNA modifications at the wobble-uridine site to regulation of viral replication.

## Results

### CHIKV infection induces a metabolic reprogramming towards one-carbon pathway

We performed an untargeted metabolomics study to uncover and compare the metabolic responses to a CHIKV infection within the first 8 h replication cycle in Vero E6 and Huh7.5.1 cells (**Fig. S1**). The 8 h time-point was chosen to avoid any confounding lytic effects usually associated with later stages of CHIKV infection^37^. Cellular metabolites were extracted with MeOH:ACN:H_2_O (2:2:1) and analyzed by liquid chromatography coupled to mass spectrometry (LC/MS) after electrospray ionization in the positive mode. On average, a total of 116 (116 from Vero E6 and 116 from Huh7.5.1), 108 (108 from Vero E6 and 108 from Huh7.5.1), and 136 (122 from Vero E6 and 151 from Huh7.5.1) metabolites were detected for multiplicities of infection (MOI) of 0.2, 2, and 20, respectively (**Table S1, Supplementary data file 1 – 3**). Applying a stringent cut-off of log_2_ fold change ≥ 1, the numbers of significantly repressed metabolites were 9, 3, 1 (Vero E6), and 5, 0, 3 (Huh7.5.1) for MOI of 0.2, 2, and 20, respectively (**Fig. 1a-f**). The numbers of metabolites which were significantly induced by CHIKV infection decreased with an increasing MOI (**Fig. 1a-f**). While relatively few metabolites were induced at MOI of 20 (2 in Huh7.5.1, and 0 in Vero E6), higher numbers were found at MOI’s of 2 (13 in Vero E6, 14 in Huh7.5.1) and 0.2 (18 in Vero E6 and 31 in Huh7.5.1).

**Fig. 1.**
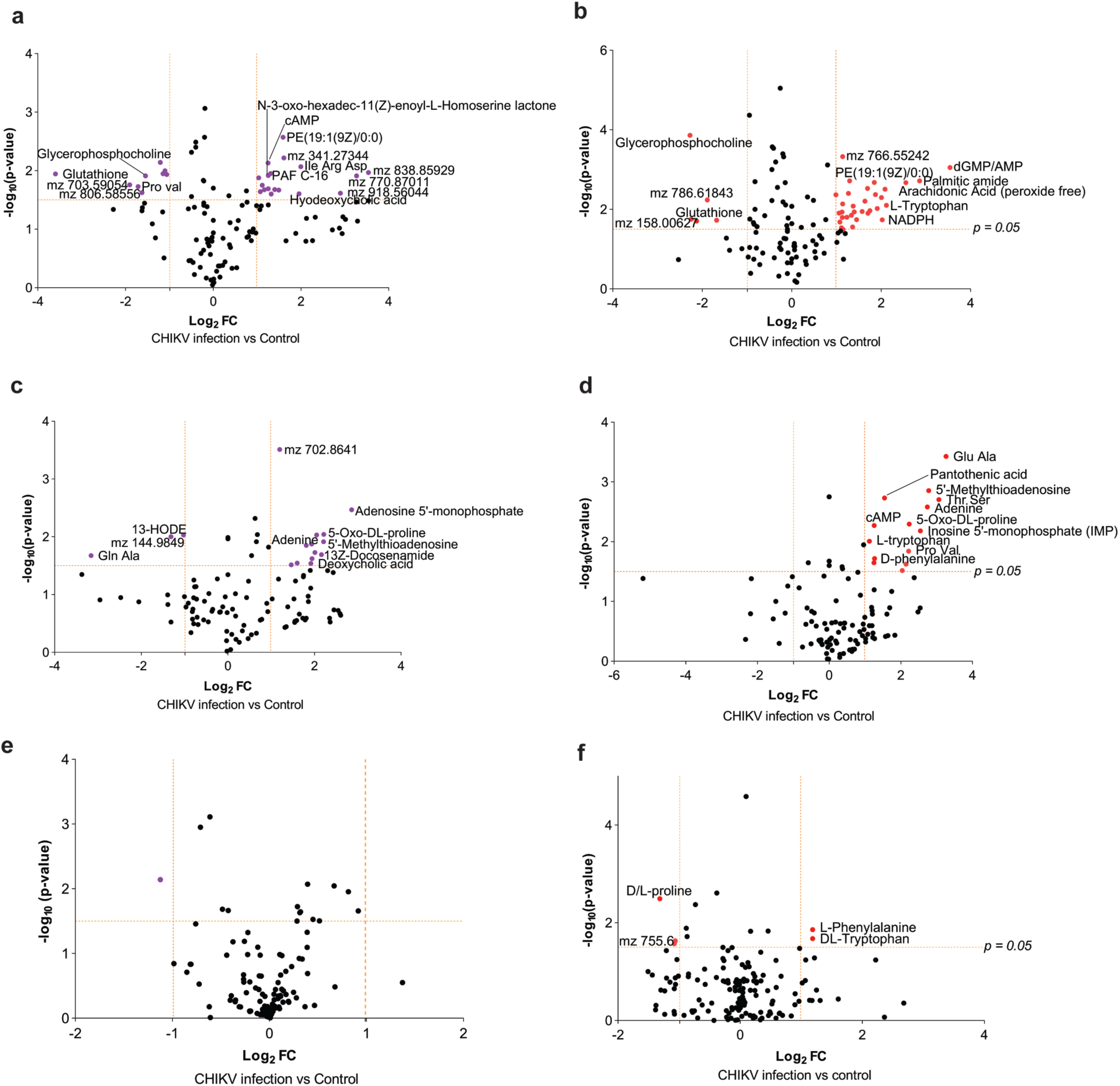
Metabolic changes induced by CHIKV infection in mammalian cells. **a-f**. Scatter plots of metabolite changes for CHIKV infection performed at the indicated multiplicity of infections (MOI). Each dot represents an average fold change (FC) of five independent biological replicates performed at multiple technical replicates (n = 5 biological replicates). Colored dots (red and pink) denote log_2_ FC ≅ 1 or –1 for the respective cell lines. Infection dose, **a** and **b**: MOI of 0.2, **c** and **d**: MOI of 2, **e** and **f**: MOI of 20, cell lines: **a**, **c**, **e**: Vero E6 and **b**, **d**, **f**: Huh7.5.1.

About 14 metabolites which were significantly induced in Huh7.5.1 cells at MOI of 2, were also induced in Vero E6 cells at MOI of 2 (**Fig. 1c-d; Table S1**). Among these were various lipid derivatives, as expected, and di– and tripeptides of which their relevance is not yet known. However, we were most attracted by the levels of MTA that changed by log_2_ 2.8-fold and 1.9-fold during CHIKV infection in Huh7.5.1 and Vero E6 cells, respectively (**Fig. 1; Table S1**). MTA is a sulfur-containing nucleoside and a by-product of polyamine biosynthesis from *S*-adenosylmethionine (SAM) (**Fig. 2a**)^38^. It is also a member of the Met salvage pathway and one-carbon (1C) metabolism, and as such important for the regeneration of methionine and single-carbon units, respectively. The first step of this Met salvage pathway is catalyzed by MTA-phosphorylase (MTAP), which leads to phosphorylation of the ribose and the cleavage of adenine. The relevance of this pathway is underlined by the observation that also adenine, adenosine-5’-monophosphate (AMP), and cyclic AMP (cAMP) were among the enriched metabolites.

**Fig. 2.**
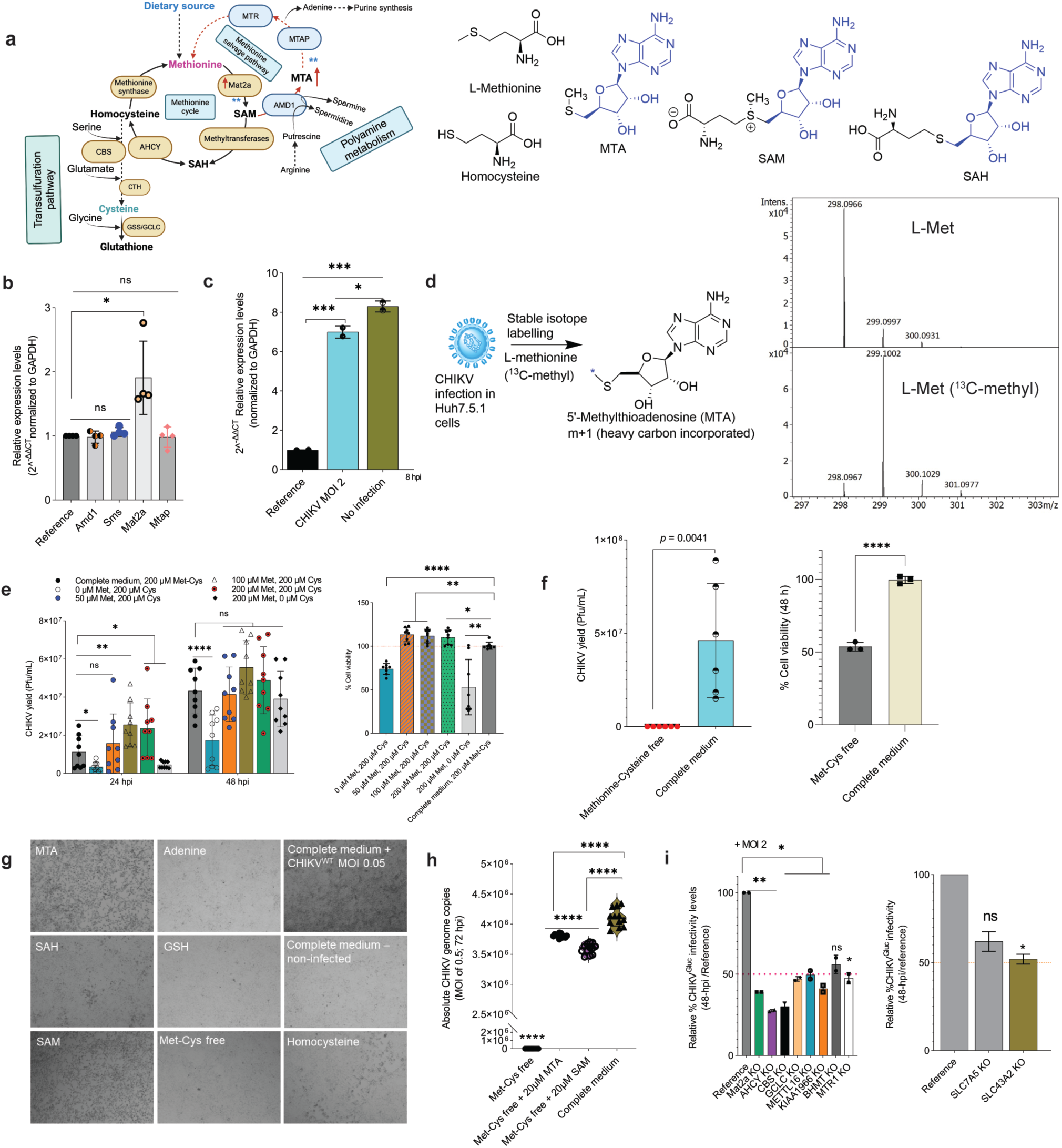
Reprogrammed Met salvage cycle through MTA induction facilitates Met – Cys availability to regulate CHIKV replication. **a**. Methionine, transsulfuration, and polyamine pathways. Abbreviations of enzymes: AHCY, *S*-adenosylhomocysteine hydrolase; AMD1, Adenosylmethionine decarboxylase 1; CBS, Cystathionine β-synthase; CTH, Cystathionine ɣ-lyase; GCLC, Glutamate-cysteine ligase; GSS, Glutathione synthase; Mat2a, Methionine adenosyltransferase 2a; MTAP, Methylthioadenosine phosphorylase; MTR, Methionine synthase. Abbreviations of metabolites: MTA, 5ʹ-methylthioadenosine; SAM, *S*-adenosylmethionine; and SAH, *S*-adenosylhomocysteine. ** and ↑ denote the CHIKV-induced upregulated enzyme (Mat2a) and metabolite (MTA). *Right*: Chemical structures of major Met cycle metabolites – Met, MTA, SAM, SAH, and homocysteine. Blue indicates structural similarity of MTA, SAM, and SAH. **b** and **c**. qPCR profiles of Met salvage cycle enzymes during CHIKV^WT^ infection at MOI of 2 for 8 hpi under complete medium (**b**) and Met-Cys limiting conditions for Mat2a expression (**c**). Experiments were conducted four times (**b**): *Student’s t*-test, *Mat2a* vs reference, t = 3.181, df = 3, * – *p* = 0.05, and twice (n = 2 independent replicates) (**c**): CHIKV MOI 2 vs reference, t = 27.29, df = 2, *** – *p* = 0.0013, No infection, t = 36.51, df = 2, *** – *p* = 0.0007, CHIKV MOI 2 vs No infection, t = 4.372, df = 2, * – *p* = 0.0485, each in three technical replicates. **d**. *Left:* Schematic of isotopic incorporation tracking of methionine during CHIKV infection (8 hpi; MOI of 2) using L-Met (^13^C-methyl) into MTA. *Right*: Mass spectra of MTA from CHIKV-infected cell cultures grown with either L-Met (top) or with L-Met (^13^C-methyl) (bottom). **e** – **f**. CHIKV replication regulation by Met-Cys availability. Infected cells were cultured in Met-Cys free or complete medium for indicated hpi and CHIKV infection intensity quantified by plaque assay (e – f). In parallel, cell viability was assessed by CellTiter-Glo^®^ assay at 48 h (e, f). *t*-test, CHIKV yield 24 hpi: Complete medium vs 0 µM Met, 200 µM Cys, t = 2.694, *p* = 0.0243, 100 µM Met, 200 Cys, t =2.995, *p* = 0.0092, 200 µM Met, 200 µM Cys, t = 2.131, *p* = 0.05, 200 µM Met, 0 Cys, t = 2.357, *p* = 0.0437. 48 hpi, Complete vs 0 Met, 200 µM Cys, t = 4.339, *p* = 0.0005, ns – not significant. **g**. Infection rescue assay assessed by supplementing the indicated sulfur-containing intermediate metabolites into Met-Cys deprived infected cells. Representative bright field microscopy images of infected cells taken 48 hpi. Scale bar, 300 µm. **h**. MTA favors CHIKV replication in a Met-Cys free medium. CHIKV infection was conducted with a wild-type virus at MOI of 0.5 for 72 hpi. Viral RNA was extracted from supernatants and absolute genome copies quantified by qPCR using standard CHIKV E1 gene fragment. *Student*’s *t*-test, Met-Cys free + SAM, t = 10.38, df = 22, *p* < 0.0001, Met-Cys free + MTA, t = 6.682, df = 22, *p* < 0.0001, Met-Cys free + MTA vs Met-Cys free + SAM, t = 8.485, df = 22, *p* < 0.0001. **i**. Comparative CRISPR deletion knockout effects of Met transporter genes and Met-Cys pathways on CHIKV infection (MOI of 2; 48 hpi). Experiments were conducted for two independent replicates in 96 technical replicates. *Student*’s *t*-test, Mat2a KO vs Reference, t = 123, df = 1, *p* = 0.0052, AHCY KO, t = 145, *p* = 0.0044, CBS KO, t = 35, *p* = 0.0182, GCLC KO, t = 53, *p* = 0.0120, METTL16 KO, t = 20.20, *p* = 0.0315, KIAA1966 KO, t = 29.50, *p* = 0.0216, MTR1 KO, t = 21, *p* = 0.0303, SLC43A2 KO, t = 24, *p* = 0.0262, ns – not significant.

### Methionine – cysteine availability orchestrated by MTA induction regulates CHIKV replication

1C metabolic pathways comprise the interlinkage of Met, folate, and transsulfuration metabolism. Previous studies have described 1C metabolism as a prime source of purines and RNA methylation through methionine (Met) and folate cycles, that is often hijacked for optimal viral RNA replication^12,14,39^. Increased production of MTA during CHIKV infection in both Vero E6 and Huh7.5.1 cells, similar to hepatitis C virus infection^40^, reflected active enzymatic activities utilizing SAM from the Met cycle (**Fig. 2a)**. Polyamines generated from the Met arm of 1C are known to support CHIKV replication^41,42^. In line with this, we evaluated the relevance of this pathway for CHIKV infectivity in more detail by exploring pharmacological inhibitions. We chose Huh7.5.1 cells for these subsequent infection analyses. The synthesis of MTA from SAM decarboxylation is catalyzed by *S*-adenosylmethionine decarboxylase (AdoMetDC or AMD1) (**Fig. 2a**). AMD1 is an important antiviral target for human immunodeficiency virus (HIV) and herpes simplex virus 1 (HSV-1)^43,44^, but nothing is known for alphaviruses. To test a possible involvement of MTA-producing polyamine enzymes for CHIKV replication, we treated CHIKV-infected cells with the specific AMD1 inhibitors mitoguazone and SAM486A and the spermine synthase (SMS) inhibitor *N*-(3-aminopropyl)cyclohexylamine^45,46^. These compounds however failed to suppress CHIKV replication up to tested concentrations of 10 µM (**Table S2**). The lack of observable inhibitory effects from these compounds implied an induction of MTA independent of polyamine metabolism under CHIKV influence. In fact, treatment with MTDIA, a potent inhibitor of cellular MTAP, that causes accumulation of MTA along the Met salvage route, could not inhibit but fairly increased CHIKV replication. Comparison of gene transcripts along the Met salvage cycle for *Mat2a*, *Amd1*, *Mtap*, and *Sms* in the infected cells depicted a significant 1.9-fold upregulation of *Mat2a*, whereas the rest had negligible changes (**Fig. 2b**), coinciding with possible CHIKV-induced sulfur depletion^47,48^. Under Met deprivation conditions, a significant 7-fold upregulation of *Mat2a* transcripts was observed in CHIKV-infected cells (**Fig. 2c**). *Mat2a* induction and increased MTA levels suggested infection-induced sulfur pool depletions prompting replenishment through the Met salvage cycle. *Mat2a* activation is reported to selectively involve 3ʹ untranslated region of MAT2a mRNA methylation writers, METTL16 and YTHDC1 (KIAA1966)^49,50^. These findings prompted us to use the stable isotope L-Met (^13^C-methyl) labelling assay to probe if CHIKV engaged the Met salvage pathway during its infection at MOI of 2 for 8 hpi (**Fig. 2d**). Feeding L-Met (^13^C-methyl) led to a mass shift for protonated MTA from 298.0966 Da to 299.1002 Da, while only a minor fraction of 8% remained unlabeled. This shows that MTA is indeed synthesized from Met under CHIKV infection conditions.

To prove the relevance of Met availability for CHIKV, we depleted Met in the culture medium before infecting cells. In culture medium and serum, the concentrations of Met is empirically estimated to be 200 µM and ≅ 30 µM, respectively^51^. To better understand the CHIKV infection dynamics on Met restriction, we first varied the Met concentration in culture medium (in absence and presence of 200 µM Cys) and quantified infection levels over 24 hpi to 48 hpi time periods by a plaque assay. Irrespective of the analysis time-point, absence of either Met or Cys negatively impacted CHIKV titres, which increased with increasing Met concentrations (**Fig. 2e**). Strikingly, deprivation of both Met and Cys for 8 hpi, 24 hpi, 48 hpi, or 72 hpi from CHIKV-infected cells completely thwarted both wild-type and reporter CHIKV replication without inducing severe collateral cytotoxicity (**Fig. 2f; Fig. S2**). Rescue of this phenotype was only possible from a weak to a greater extent upon exogenous addition of sulfur-containing intermediates such as 100 µM homocysteine, 20 µM SAM, and 20 µM MTA. However, the addition of 20 µM glutathione (GSH), 20 µM adenine, or 20 µM SAH to Met-Cys free medium had no effect (**Fig. 2g**). Moreover, stable isotope tracing of L-Cys-^13^C_3_, ^15^N during CHIKV infection did not lead to a notable mass shift of cellular metabolites (**Supplementary data file 4**). These findings suggested a combined Met-Cys antiviral effect, as pharmacological inhibition of *de novo* Cys biosynthetic enzymes from Met showed no antiviral effect (**Table S2**) and deprivation of either sulfur amino acid was not sufficient to clear CHIKV infection (**Fig. 2e**). Moreover, the induced MTA during CHIKV infection proved to be a directly pro-viral metabolite and was utilized upon sulfur pool depletion, as exogenously added MTA compensated for the lack of Met-Cys and act as an alternative fuel for CHIKV replication more effectively than SAM (**Fig. 2h**).

Uptake of Met and Cys from the extracellular environment is known to be regulated by transmembrane solute transporter channels of L-type amino acid transporters (LAT) 1 to 4 (SLC7A5, SLC7A8, SLC43A1, and SLC43A2, respectively) and xCT (SLC7A11) systems^52^. We tested if blockade of these channels with specific inhibitors would recapitulate Met-Cys deprivation effects. Infected cells were treated with the LAT inhibitors 2-aminobicyclo-(2,2,1)-heptane-2-carboxylic acid (BCH) and JPH203, and the xCT inhibitors sulfasalazine and erastin. Contrary to our expectation, none of these compounds exerted a meaningful anti-CHIKV activity (**Table S2**). We confirmed through a qRT-PCR that in the absence of Met-Cys, expression of SLC7A5, SLC43A2, and SLC7A11 genes was upregulated > 3-fold in CHIKV-infected cells (**Fig. S3a**). We therefore decided to interrogate if SLC7A5 and SLC43A2 deletion could abrogate CHIKV infection under Met-Cys rich medium conditions. In parallel, the dependency of CHIKV infection on Met-Cys-involving pathway enzymes was also probed by CRISPR/Cas9-mediated deletions. Knockouts of Mat2a and its regulators (METTL16 and KIAA1966), *S*-adenosylhomocysteine hydrolase (AHCY), methyltransferase ribozyme 1 (MTR1), betaine-homocysteine methyltransferase (BHMT), cystathionine β-synthase (CBS), and glutamate-cysteine ligase (GCLC) were generated. Transporter channel deletion failed to disclose a clear link between the dependency of Met uptake and viral infectivity as observed with Met-Cys deprivation, despite a ∼40-50% infectivity reduction (**Fig. 2i**). Consistent with the upregulation of Mat2a, its knockout together with that of the regulator YTHDC1 (KIAA1966), but not METTL16, equally led to a 60% CHIKV infectivity reduction (**Fig. 2i**). Stronger antiviral effects were exerted upon deleting AHCY (72%) and CBS (70%) (**Fig. 2i**), implicating that Cys metabolism is essential to complete CHIKV replication. Collectively these data indicate that CHIKV uses MTA and the activity of Mat2a for sulfur resupply to favor its replication.

### ALKBH8 impairment associated with tRNA hypomodification recapitulates Met-Cys deprivation anti-CHIKV effects

Sulfur amino acids are also required during the maturation process of transfer RNA (tRNA) by facilitating sulfur-relay pathway thio-modifications for an enhanced translational fidelity and structural stability (**Fig. S4**)^48^. We reasoned that CHIKV could maximize its RNA replication by mobilizing host cellular sulfur pools engaging either through stress-mediated tRNA modifications at the anticodon wobble site (uridine 34; U_34_), or thiol-disulfide mediated exchanges^34,53^. To test this link, we performed a comparative gene expression profiling to define the responsive players of tRNA modifications upon Met-Cys deprivation in CHIKV-infected cells. **Fig. 3a** depicts that CHIKV infection induced 2.1-, 1.8-, and 1.3-fold changes in the expression of *Kiaa1456*, *Ctu1*, and *Urm1*, respectively, in complete medium. Under Met-Cys deprivation conditions, infected cells upregulated transcripts for *Kiaa1456*, *Alkbh8*, and *Mocs3* by ≥ 4-fold, and those for *Ctu2* and *Nfs1* increased by ≥ 2-fold, suggesting an active interaction with tRNA modifiers. In contrast, *Ctu1* levels were remarkably reduced by 2.25-fold relative to complete medium condition while *Elp3* and *Urm1* appeared steadily responsive to CHIKV infection regardless of the nutrient status. We then explored the relative effects of these tRNA modifying proteins on CHIKV infection in complete medium using CRISPR knockouts. The results revealed that ALKBH8 and Urm1 knockouts profoundly restricted CHIKV infectivity to 28% and 35%, respectively (**Fig. 3b**). A knockout of ALKBH8 did not affect cell viability (**Fig. 3b**). ALKBH8 is a paralog of human KIAA1456, and specifically catalyzes the last methylation step to give modified 5-methylcarbonylmethyluridine (mcm^5^U) providing the substrate for the U_34_-tRNA 2-thiolation reactions to 5-methoxycarbonylmethyl-2-thiouridine (mcm^5^s^2^U) (**Fig. 3c, S4**)^54–57^. Whilst mcm^5^s^2^-tRNA modifications are described to be translationally silenced during CHIKV infection^34^, current thiol deprivation responses and the strong infection restriction observed upon of ALKBH8 KO (**Fig. 3a-b**) suggested a link to sulfur availability for tRNA modification required for CHIKV replication. We further explored if supplementing mcm^5^U or mcm^5^s^2^U into ALKBH8 KO cells would rescue CHIKV infectivity. The results inferred that exogenous addition of mcm^5^s^2^U, but not of mcm^5^U, led to a significant 1.24 ± 0.06-fold increment in CHIKV infection regardless of tested concentrations and assay timing (**Fig. 3d**). Moreover, comparative mass spectrometry quantification of U_34_-tRNA modifications confirmed that cells from both ALKBH8 KO and Met-Cys free conditions had significantly reduced levels of mcm^5^U, but only ALKBH8 KO exhibited significantly reduced mcm^5^s^2^U levels (**Fig. 3e**). These findings confirmed that ALKBH8 perturbation was detrimental to CHIKV, recapitulating Met-Cys deprivation effects, and infectivity rescue was slightly possible on addition of thiol-containing mcm^5^s^2^U.

**Fig. 3.**
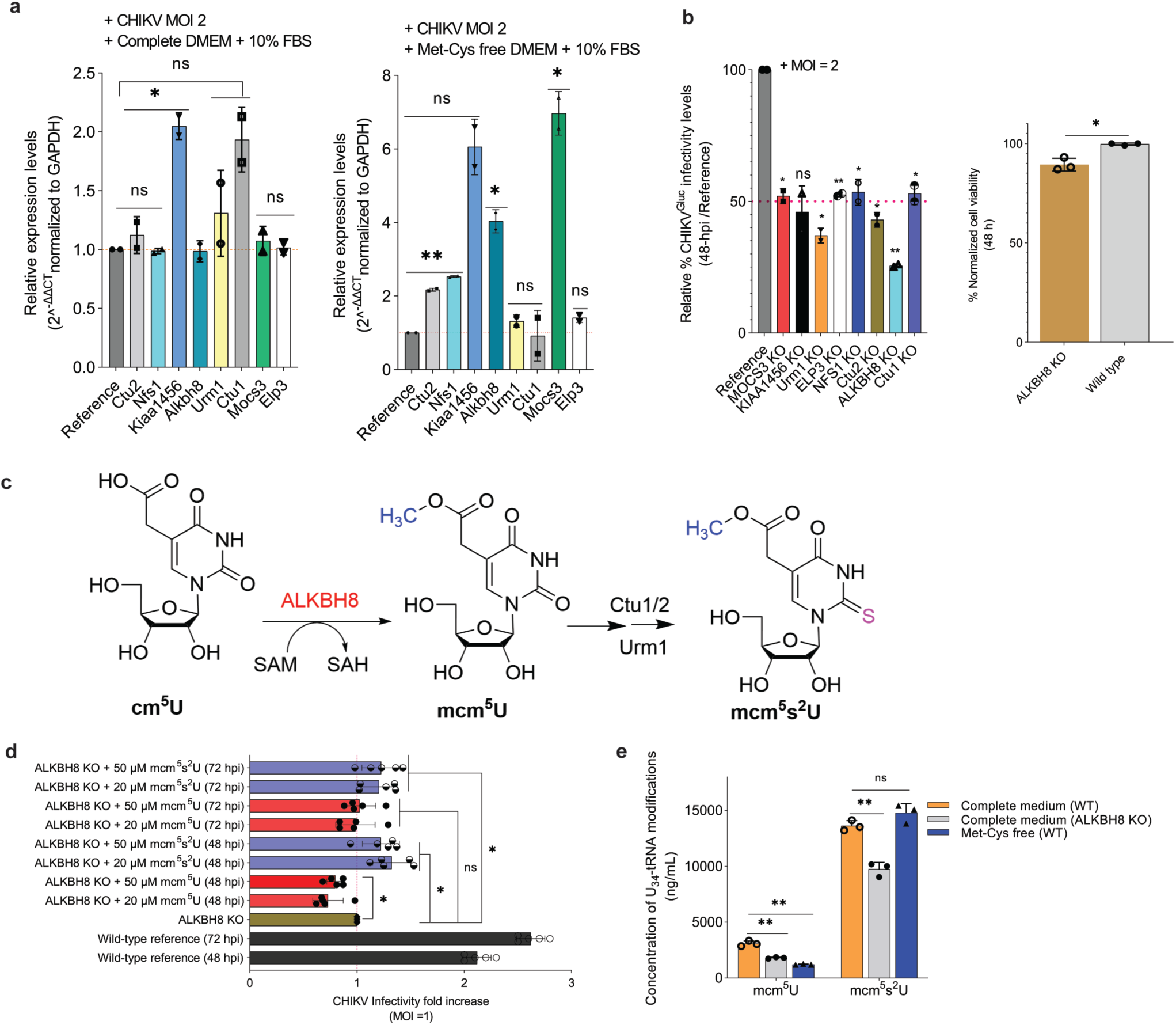
ALKBH8 deletion recapitulates Met-Cys deprivation effects on CHIKV infectivity. **a.** Expression levels of tRNA modifying enzymes during CHIKV^WT^ infection at MOI of 2 at 8 hpi under complete medium and Met-Cys depleted conditions, determined by qPCR. Experiments were conducted twice (n = 2 independent replicates) in three technical replicates. *Student*’s *t*-test: Complete medium, *Kiaa1456* v reference, t = 13.28, df = 1, *p* = 0.0478. Met-Cys free medium: *Ctu2*, t = 41.86, *p* = 0.0152, *Nfs1*, t = 76.5, *p* = 0.0083, *Alkbh8*, t = 13.49, *p* = 0.0471, *Mocs3*, t = 14.21, *p* = 0.0447, ns – not significant. **b**. *Left*: Comparative CRISPR deletion effects of tRNA modifying genes on CHIKV infection (MOI of 2; 48 hpi). Experiments were conducted for two independent replicates (96 technical replicates). *Student*’s *t*-test, MOCS3 KO, t = 24, df = 1, *p* = 0.0265, Urm1 KO, t = 31.50, df = 1, *p* = 0.0202, ELP3 KO, t = 95, df = 1, *p* = 0.0067, NFS1 KO, t = 13.29, df = 1, *p* = 0.0478, Ctu2 KO, t = 28.50, df = 1, *p* = 0.0223, ALKBH8 KO, t = 149, df = 1, p = 0.0043, Ctu1 KO, t = 15.67, df = 1, *p* = 0.0406, ns – not significant. *Right*: ALKBH8 KO cell viability tested at 48 h. *Student*’s *t*-test, t = 5.480, df = 4, * – *p* = 0.0276. **c**. Reaction scheme of ALKBH8-dependent U_34_-tRNA modifications. Blue and pink atoms indicate the specific methylation and thiolation sites. **d**. Comparative effects of exogenous supplementation of mcm^5^U and mcm^5^s^2^U to ALKBH8 KO cells on CHIKV infectivity for 48 and 72 hpi (MOI of 1; n = 5). *Student*’s *t*-test, 48 hpi, ALKBH8 KO + 20 µM mcm^5^U, t = 4.227, df = 4, *p* = 0.0134, ALKBH8 KO + 50 µM mcm^5^U, t = 5.6154, df = 4, *p* = 0.005, ALKBH8 KO + 20 µM mcm^5^s^2^U, t = 4.0833, df = 4, *p* = 0.0151, ALKBH8 KO + 50 µM mcm^5^s^2^U, t = 2.9074, df = 4, *p* = 0.0438. 72 hpi, ALKBH8 KO + 20 µM mcm^5^s^2^U, t = 2.7271, df = 4, *p* = 0.05, ALKBH8 KO + 50 µM mcm^5^s^2^U, t = 2.8105, df = 4, *p* = 0.0482, ns – not significant. **e**. Quantification of mcm^5^U and mcm^5^s^2^U tRNA modifications by LC-MS. Two-tailed *Student*’s *t-*test, mcm^5^U: Complete medium (WT) vs Met-Cys free (WT), t = 13.12, df = 3, ** – *p* = 0.0048, Complete medium (WT) vs Complete medium (ALKBH8 KO), t = 8.812, df = 3, ** – *p* = 0.0083. mcm^5^s^2^U: Complete medium (WT) vs Met-Cys free (WT), t = 2.079, df = 3, ns – *p* = 0.1259, Complete medium (WT) vs Complete medium (ALKBH8 KO), t = 8.695, df = 3, ** – *p* = 0.0013.

We next analyzed the relationship between ALKBH8 and the pro-viral Mat2a expression and MTA increment under CHIKV infection conditions. We established that, unlike MTA and cycloleucine treatments (Mat2a inhibitor), CHIKV infection of ALKBH8 KO or Mat2a KO cells under normal culture conditions did not trigger upregulation of either gene. These conditions rather resulted in no effect on Mat2a expression and downregulation of ALKBH8 expression, respectively (**Fig. S3a**), suggesting an impaired crosstalk. In contrast to this observation, the pharmacological inhibition of Mat2a by 30 mM cycloleucine caused an upregulation of ALKBH8, and Mat2a levels increased, when 20 µM MTA was added. Together, these data confirm ALKBH8 as a relevant tRNA modifier that fine-tunes CHIKV infection under the control of sulfur availability.

### Exogenously-added MTA exerts m^6^A-independent priming of CHIKV replication

Exogenous uptake of MTA sufficiently supported CHIKV replication in absence of Met-Cys or SAM. To illuminate the underlying mechanism, at first a Met-Cys time-of-deprivation assay was conducted for a single CHIKV replication cycle (MOI of 1). The most clearly pronounced inhibitory effect was indeed observed during 0 – 2 h post-entry, followed by a general viral suppression plateau for the other lifecycle steps (**Fig. 4a**). This inhibitory time-window corresponds to the period of post-transcriptional modifications and translation of CHIKV RNA preceding viral genome replication^36,37^. We therefore explored the relative priming effects of MTA and its impact on CHIKV RNA. For other viruses it is known that methylation of adenosine to N^6^-methyladenosine (m^6^A) stably initiates viral RNA processing^36,58,59^. As the metabolite MTA is related to intracellular methylation precursors through Met metabolism, we evaluated if MTA influences the methylation status of the RNA base m^6^A under Met-Cys free infection conditions. Using m^6^A RNA-immunoprecipitation followed by fluorescence quantification, we observed a clear reduction of m^6^A levels upon Met-Cys deprivation. However, the exogenous supply of MTA significantly failed to increase the RNA methylation signal to the level of the infection reference (**Fig. 4b**), suggesting an alternative priming mode.

**Fig. 4.**
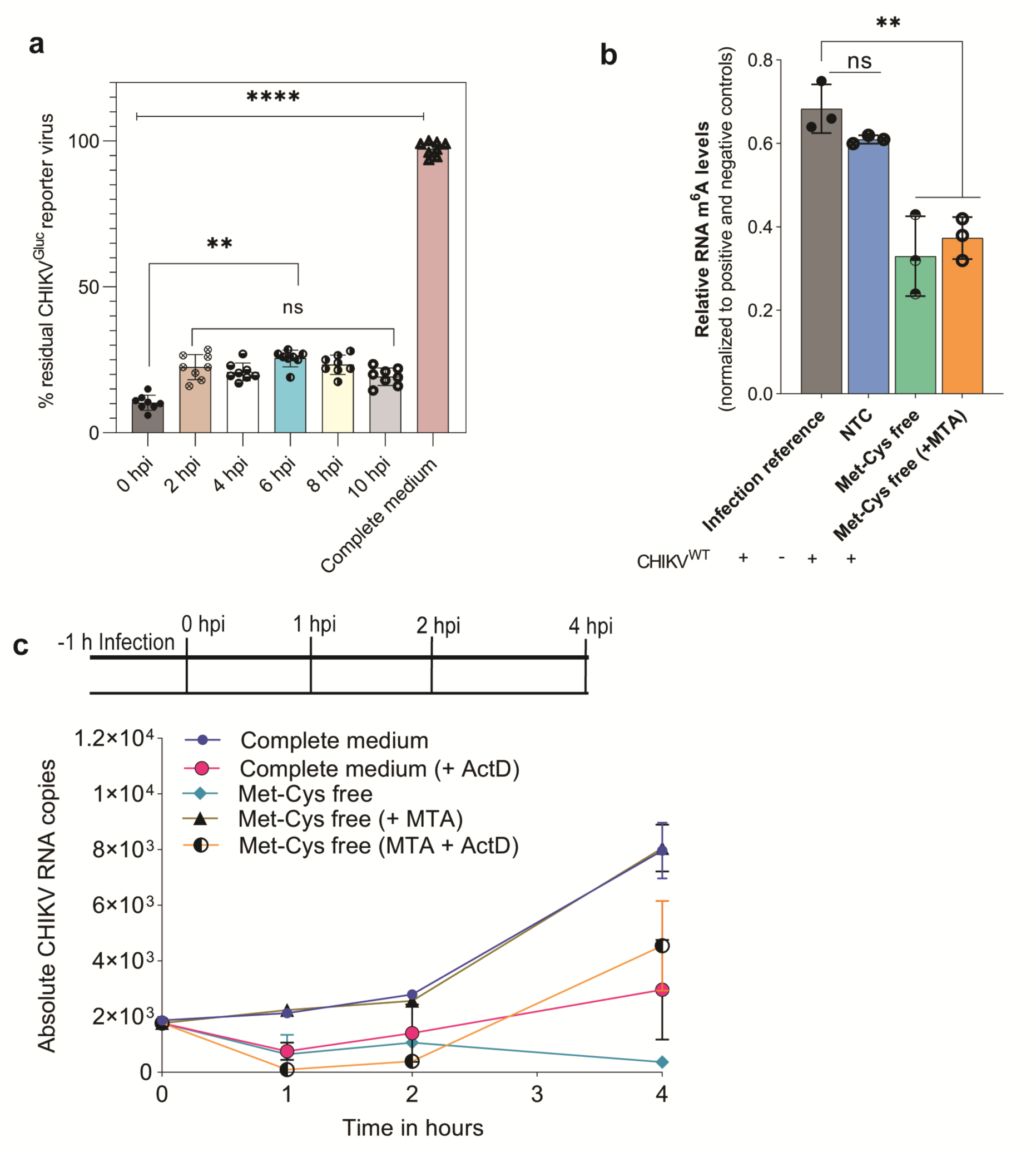
MTA priming effects on CHIKV RNA replication. **a**. Met-Cys time-of-deprivation assay at MOI of 1 for a single replication cycle. Infection was initiated with complete culture medium and Met-Cys free medium was added at the indicated time and viral infection intensity quantified at 24 hpi. **** – *p* < 0.0001, ** – < 0.05, and ns, not significant (*Student’s t*-test). **b**. Relative m^6^A RNA methylation analysis using EpiQuik m^6^A RNA methylation quantification fluorescence kit. Unpaired two-tailed *Student*’s *t*-test; NTC vs Infection reference, t = 2.137, df = 4, ns – *p* = 0.0994, Met-Cys free vs Infection reference, t = 5.467, df = 4, ** – *p* = 0.0054, Met-Cys free + MTA vs Infection reference, t = 6.951, df = 4, ** – *p* = 0.0024. **c**. Analysis of CHIKV RNA transcription priming by MTA. Total RNA samples were collected from MTA-supplemented Met-Cys free + CHIKV-infected cells in presence of 10 µM Actinomycin D (ActD) or its absence (complete medium) at the indicated time points post cell entry. CHIKV RNA copies were quantified by qRT-PCR analysis of E1 gene. Experiments were performed in triplicates for three replicates (n = 3).

Relooking into the responsive profile of CHIKV to the structurally-related MTA and SAM (**Fig. 2a, h**), the possibility to prime viral RNA transcription processing in a relatively similar fashion was hypothesized as previously described^60,61^. To elucidate this possibility, infected cells were assessed for CHIKV RNA genome copies during the transcription phase in presence or absence of 10 µM actinomycin D (ActD), an inhibitor targeting DNA-dependent RNA polymerases. Compared to infected cells in complete medium or in Met-Cys free medium supplemented with MTA, the addition of ActD reduced the viral RNA genome copies at least during the first two hours of post-entry (**Fig. 4c**). However, this ActD-mediated arrest was only transient in the Met-Cys free medium supplemented with MTA as CHIKV RNA levels increased between 2 hpi and 4 hpi during viral RNA synthesis phase. The early ActD mediated inhibition of MTA-mediated pro-viral effects strongly suggests that the MTA priming effect on CHIKV is primarily exerted at pre-transcribed mRNA complexes. These data provide an important insight to demonstrate that the pro-viral MTA further plays an essential role during viral RNA transcriptional-replication events to license CHIKV RNA replication independent of cellular m^6^A RNA status.

### Deazaneplanocin A and Adenosine dialdehyde are potent CHIKV inhibitors

Finally, we tested if the CHIKV dependency on sulfur metabolism could be explored for a therapeutic approach with known pharmacological small-molecules. CHIKV-infected cells were treated with known inhibitors of MAT2a (FIDAS-3, AG-270, PF-9366), AHCY (3-deazaneplanocin A HCl (DzNep) and adenosine dialdehyde (Adox)), Mtr1 (tubercidin and trifluoromethyltubercidin (TFMT)), and 1C metabolism enzymes (**Table S2**). From this antiviral screen, only the AHCY inhibitors, DzNep and Adox potently inhibited CHIKV replication with EC_50’s_ of 0.42 ± 0.05 nM and 14.0 ± 0.4 nM, respectively, in the CHIKV^Gluc^ assay (**Fig. 5a-c**). The high activities were confirmed in the wild-type CHIKV cytopathic assay, with EC_50_’s of 1.7 nM and 139.9 nM, respectively (**Fig. S5**). AHCY represents a less explored alphaviral target to block pro-viral transmethylation reactions^62,63^. DzNep had previously been reported not to show a direct inhibition of CHIKV nsP1 MTase activity^64^, so that our antiviral activity data confirm an exploitation of a host cellular target with a high anti-CHIKV^Gluc^ activity. AHCY inhibitors are also capable of destabilizing histone methylation by indirectly inhibiting enhancer of zeste homolog (EZH) proteins^65^. To rule out such a possibility from specific CHIKV inhibition, other EZH1/2 and DNA methyltransferase (DNTM) inhibitors were tested. Only EZH2-targeting PF-06726304 exerted anti-CHIKV activity with an EC_50_ of 1.13 ± 0.1 µM. All the other tested inhibitors were inactive (**Table S2**). This finding suggests that the observed anti-CHIKV effect of AHCY inhibitors was more due to methionine cycle than to EZH1/2 homolog inhibitions, and their effects to a greater extent phenocopied Met-Cys deprivation at 8 hpi (**Fig. 5d-e**). We then addressed the question why Mat2a inhibitors were inefficient at suppressing CHIKV replication, whereas *Mat2a* deletion did. AG-270 and PF-9366 are allosterically-acting compounds, while FIDAS-3 is an unnatural substrate inhibitor. To address this discrepancy, we used the known substrate inhibitor cycloleucine at 30 mM concentration as described^49,50^ to suppress Mat2a activity. A ∼ 75% reduction in CHIKV infectivity was achieved (**Fig. 5f**), phenocopying deletion knockouts and re-affirming Mat2a as a viable CHIKV antiviral target. This observation could suggest a strong preference for substrate competitive inhibitors, though we are aware that the high cycloleucine concentration of 30 mM could also lead to secondary non-target effects.

**Fig. 5.**
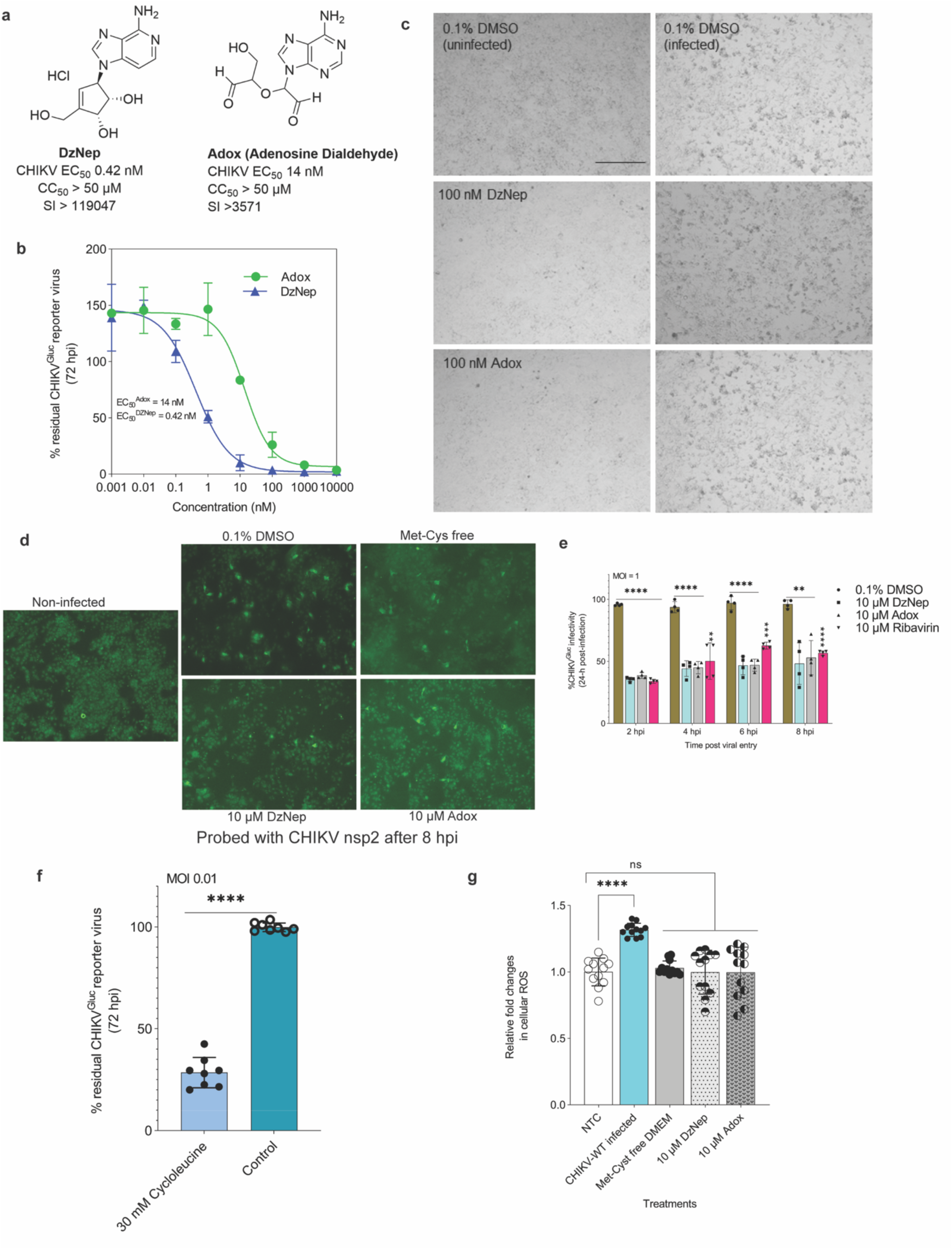
AHCY inhibitors are potent CHIKV inhibitors. **a.** Chemical structures of AHCY inhibitors; 3-deazaneplanocin A HCl (DzNep) and adenosine dialdehyde (Adox) exerting CHIKV antiviral activities at indicated potencies. **b.** Dose-response curves of antiviral effects exerted on CHIKV^Gluc^ MOI of 0.01 by DzNep and Adox for 72 hpi (n = 6 independent replicates). **c**. Representative images of bright field microscopy of infected cells (MOI of 0.01) on treatment with DzNep and Adox taken at 72 hpi. Scale bar, 300 µm. **d**. Immunofluorescence image of treatment effects of DzNep, Adox, and Met-Cys free medium on CHIKV infection for 8 hpi at MOI of 2. Infection intensity was probed using CHIKV nsP2 antibody under GFP background. **e**. Time-of-addition assay for DzNep and Adox for 8 hpi, with reference to ribavirin. Treatment was started following virus cell entry and quantification performed at 24 hpi. Assay conducted in triplicates at MOI of 1 for two independent replicates. *Student’*s *t*-test, **** – *p* < 0.0001, ** – 4 hpi: Ribavirin, t = 5.6493, *p* = 0.007, *** – 6 hpi: t = 11.377, *p* = 0.0002, ** – 8 hpi: DzNep, t = 5.5941, *p* = 0.0091, Adox, t = 6.0724, *p* = 0.0064. **f**. Cycloleucine antiviral activity at 30 mM against a non-treated control. Unpaired *Student*’s *t*-test, t = 25.99, df = 8, ****-*p* < 0.0001. **g**. Met-Cys deprivation and DzNep/Adox treatments restores CHIKV-induced reactive oxidative stress (ROS) levels. Cells were infected in complete medium, when the inhibitors were added. NTC, non-infected treatment control. *Student’*s *t*-test, CHIKV WT vs NTC, t = 9.494, df = 16, ****-*p* < 0.0001, Met-Cys free vs NTC, t = 0.8610, df = 16, *p* = 0.4017, DzNep vs NTC, t = 0.0296, df = 16, *p* = 0.9767, Adox vs NTC, t = 0.0531, df = 16, *p* = 0.9582. *ns* – not significant.

Reactive oxygen species (ROS) arising from redox imbalance are associated with CHIKV infection to facilitate autophagosome formation^66^. At 8 hpi, a significant 1.3-fold increase in cellular ROS levels was induced (**Fig. 5g**), and two key cellular biomarkers of stress-associated redox reactions, GSH and NADPH, were eminent (**Table S1**). Treatment of infected cells with the GSH synthesis inhibitors buthionine sulfoximine (BSO) and aminooxyacetic acid hemihydrochloride (OAA) failed to suppress CHIKV infection (**Table S2**). This observation implied that ROS production was just an infection epiphenomenon as previously reported for a closely related alphavirus Mayaro virus (MAYV)^67^. A reversal effect of the CHIKV-induced ROS levels was exerted on treatment of infected cells with DzNep, Adox, or Met-Cys free medium (**Fig. 5g**). In summary, these data confirm that, while Mat2a is pro-viral for CHIKV, the specific inhibition of viral Met-Cys dependency was more successful through treatment with AHCY inhibitors resulting in nanomolar antiviral effects.

## Discussion

Viral infections lead to perturbations of host cell metabolism, which were not yet comprehensively studied for CHIKV infections. Using untargeted metabolomics, we observed that CHIKV infection reprograms Met metabolism, elevating the levels of the methionine salvage cycle intermediate MTA, and also of the MTA-degradation product adenine. Gene expression studies showed the upregulation of Mat2a, which is the enzyme leading to SAM formation from Met and also its sensor, with SAM being the metabolic precursor of MTA. Further experiments revealed a strong regulation of viral replication by the availability of Met and Cys (Met-Cys) which constitute the major cellular sulfur pool suppliers. A key finding was that CHIKV replication could be primed by MTA of in absence of the Met-Cys, that implied the involvement of MTA and Mat2a to facilitate sulfur supply during CHIKV infection. This finding was further supported by associating tRNA modifications specifically catalyzed by ALKBH8 with Met-Cys deprivation effects. Finally, we identified that AHCY inhibitors were nanomolar anti-CHIKV agents and additionally acted to potently reverse virus-induced oxidative stress. **Fig. 6** illustrates our working model.

**Fig. 6.**
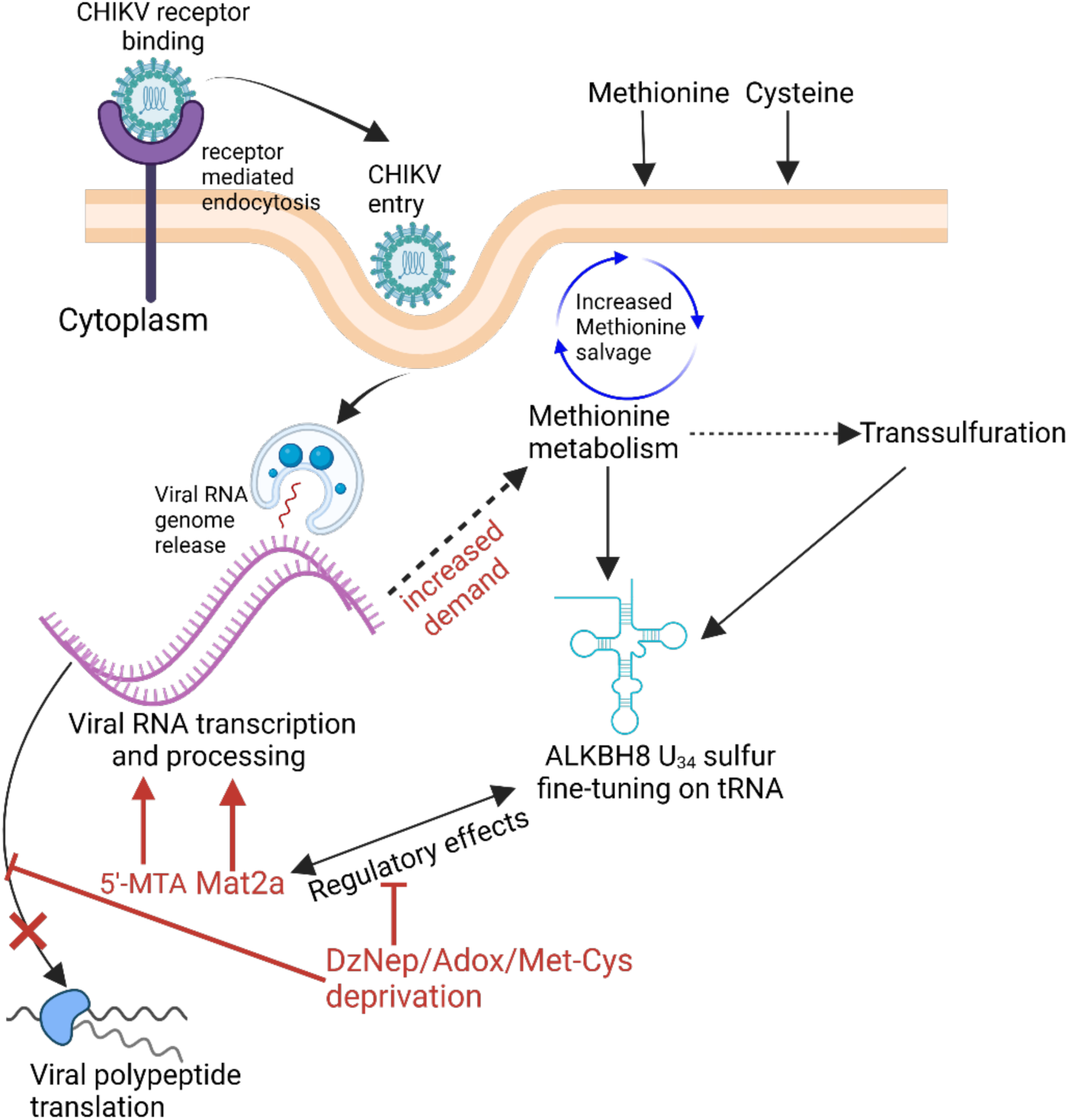
Schematic of CHIKV replication and its dependency on sulfur-associated pathways. CHIKV entry into host cell increases demand for metabolite supply from the methionine pathway, reprogramming the Met salvage pathway into increased production of 5ʹ-MTA, and upregulation of Mat2a. The reprogramming is tightly linked to the availability of sulfur amino acids that maintain downstream transsulfuration pathway to fine-tune CHIKV-modulated ALKBH8 activity. MTA can enhance CHIKV replication in absence of sulfur. Sulfur-dependent processes are targetable to thwart viral infection using the AHCY inhibitors; DzNep and Adox that mimic sulfur deprivation effects. Graphic created on Biorender.com.

Recognition of metabolic reprogramming of cellular 1C pathways to favor virus replication is well-documented for SARS-CoV-2^14^ and latency-associated human viruses^68,69^, potentially via the supply of purines and maintenance of methylation status. Elevation of MTA levels relative to its polyamine counterparts^42,70–72^ has rarely been associated with human viral infections. Until now MTA is mainly considered a potentially toxic oncometabolite that accumulates in liver cells lacking MTAP activity^73^. Only hepatitis C virus (HCV) has been reported to dysregulate MTA levels, which was interpreted to participate in alleviating virus-induced oxidant and inflammatory effects^40^. MTA increases in CHIKV-infected cells from both human liver (Huh7.5.1) and African Monkey kidney (Vero E6), concomitant with unchanged *Mtap*, *Amd1*, and *Sms* transcript levels, clearly depicted CHIKV-specific inductions as further demonstrated by stable isotope labelling. While such effects of MTA could not be exempted from viral-induced stress, the elevated levels could also have resulted from an increased activity of the induced Mat2a, which occurs as a response to sulfur depletion^47,49,50^, leading to the observation that CHIKV infection is predominantly driven by Met-Cys availability. Indeed, Mat2a is reported to be a pro-viral factor for a number of RNA viruses^32,33,74^. Interestingly, none of previous studies has associated either Mat2a or MTA with CHIKV infection. Insufficient availabilities of the key sulfur amino acids during CHIKV infection were the key driver of the observed induction of Mat2a and MTA, and not the virus-induced transmembrane uptake transporters, SLC7A5 and SLC43A2. We could further delineate the importance of the sulfur amino acids for CHIKV infection by genetic ablations of Mat2a and its regulator KIAA1966 (YTHDC1), and of AHCY and CBS, which are involved in cysteine biosynthesis from the methionine cycle. Knockout of these genes resulted in 60% to 72% reduction of the virus and confirmed the relevance of this cycle for CHIKV infection. AHCY has been demonstrated to be a pro-viral protein for ZIKV by facilitating N^6^ adenosine methylation^12^.

The dependency of viral RNA replication on sulfur amino acids could be traced to tRNA epitranscriptome dynamics^75–77^. ALKBH8 is a tRNA methyltransferase and, hence, one of the important components of epitranscriptome regulation. We had observed an upregulation of its transcription by CHIKV only in absence of the sulfur sources. Its deletion recapitulated Met-Cys deprivation effects that impaired viral replication. This antiviral effect was associated with decreased levels of mcm^5^U and mcm^5^s^2^U tRNA modifications, which rely on both ALKBH8-catalyzed methylation and the subsequent thiolation^55^. Another possibility is that the interventional deletion of ALKBH8 could restrict viral-induced polypeptide translation efficiency for which bypassing deletion effects with exogenous mcm^5^U could not rescue the infection phenotype, but the thiol mcm^5^s^2^U could at least to some extent. Preceding data had reported a favored requirement for KIAA1456, a paralog of ALKBH8, to promote CHIKV and not ALKBH8-catalyzed associated thiolations^34^. However, our data point to the cooperative nature of mcm^5^U and mcm^5^s^2^U during CHIKV infections.

From our data we conclude that the relevance of the metabolic intermediate MTA in CHIKV infection is beyond a role in metabolism, but that it also favorably primes CHIKV RNA replication under sulfur depletion. If MTA incorporation into Met salvage pathway re-initiates SAM synthesis from Met in Met-Cys deprived medium, the m^6^A signal could have increased by MTA supply almost to the level of m^6^A RNA methylation of infected cells in complete medium. However, the relative m^6^A RNA methylation status of cells, which were in infected in Met-Cys deprived medium was not increased by MTA. This indicated a different mechanism from the canonical m^6^A-mediated viral RNA replication. m^6^A post-transcriptional RNA modification is a debated aspect of arboviral RNA genomes, including CHIKV^36,78^. Lack of a strong RNA methylation effect from the addition of MTA prompted us to suggest that it also could directly act as a riboswitch ligand in a similar fashion as SAM to facilitate CHIKV RNA processing, although further experiments are needed to confirm this hypothesis. Metabolite-binding riboswitches have been already studied^79–81^ and demonstrated to exist on the structural elements of non-coding regions in viral RNA^82,83^ or engineered to conditionally control specific virus gene expressions^84–86^. The proximate binding of MTA onto CHIKV RNA structural elements^87^ could induce a stable conformation or form a non-canonical 5ʹ-end viral metabolite-capped RNA, similar to HCV^88^, human cytomegalovirus (HCMV)^89^, and DENV^90^, which is sufficient to license transcriptional initiation few hours post-entry.

Antiviral therapy by targeting host factors is a promising strategy for the discovery of potent anti-CHIKV agents^91^. Antiviral agents of this nature with nanomolar potency are rare, and the most potent agent described so far is the benzoannulene CHIKV inhibitor 8q (*N*-(4-(2-cyanopropan-2-yl)phenyl)-3-methoxy-6,7,8,9-tetrahydro-5*H*-benzo[7]annulene-2-carboxamide) targeting the human dihydroorotate dehydrogenase (DHODH) with an EC_90_ value of 270 nM^92^. This inhibitor limits the nucleotide pools available to CHIKV replication machinery. We now identified two highly potent AHCY inhibitors, DzNep and Adox, which inhibit CHIKV replication in the lower nanomolar range (EC_50s_ = 0.42 nM and 14 nM, respectively) and with a better efficacy than the related AHCY inhibitor, 6ʹ-fluoroaristeromycin analog 2c (EC_50_ = 0.13 µM)^93^. The antiviral effects of DzNep have been described for SARS-CoV-2^33,94^, Influenza Virus^95^, ZIKV^12^, Yellow Fever Virus, Vesicular Stomatitis Virus, Parainfluenza type 3 Virus, and Vaccinia viruses^96^. Such broad-spectrum activity was mediated through inhibition of viral RNA transcription and translation initiation by impairing the recruitment of the RNA-binding protein hnRNPA1 to m^6^A-modified viral RNA in addition to the modulation of antiviral immune responses. Both DzNep and Adox are pan-methyltransferase inhibitors through AHCY blockade, and thwarted CHIKV RNA replication by strongly antagonizing the pro-viral effects of MTA and Mat2a.

In conclusion, an unprecedented metabolic link involving MTA and Mat2a to sulfur homeostasis during CHIKV infection was uncovered to drive viral RNA processing. The linkage of Mat2a upregulation to Met-Cys deprivation was traced to U_34_-tRNA modifications and implicated ALKBH8 into the infection metabolism complex of CHIKV. Importantly, the identification of AHCY inhibitors as potent anti-CHIKV agents should reinvigorate an accelerated development of host-targeting therapies tailored to the identified pro-CHIKV metabolic pathways and its associated downstream vulnerable targets.

## Methods

### Materials

*S*-adenosylmethionine (SAM, Cat# A4377), *S*-adenosylhomocysteine (SAH, Cat# A9384), glutathione (Cat# G-4251), Actinomycin D (Cat# A9415), bovine serum albumin (Cat# A9418), DL-Homocysteine (Cat# 44925), Cycloleucine (Cat# A48105), puromycin (Cat# P4512), QC1 (Cat# SML0481), and UNC1999 (Cat# SML0778) were obtained from Sigma-Aldrich. JPH203 (Cat# Cay29715), BCH (Cat# Cay15249), Sulfasalazine (Cat# Cay15025), MTDIA (Cat# Cay29711), SAM486A (Cat# Cay34901), PF-9366 (Cat# Cay30782), LY345899 (Cat# Cay36627), DS44960156 (Cat# Cay28360), CPI-169 (Cat# Cay18299), and 5-azacytidine (Cat# Cay11164) were purchased from Cayman Chemicals. FIDAS-3 (Cat# E0081), Adenosine dialdehyde (Adox, Cat# S8608), Gambogenic acid (Cat# S9031), GSK126 (Cat# S7061), GSK343 (Cat# S7164), and PF-06726304 (Cat# S8494) were purchased from Selleck Chemicals. SHIN-1 (Cat# HY-112066), Wedelolactone (Cat# HY-N0551), DS18561882 (Cat# HY-130251), Erastin (Cat# HY-15763), AG-270 (Cat# HY-138630), Mitoguazone (Cat# HY-106634), Tubercidin (Cat# HY-100126/CS-5578), Trifluoromethyltubercidin (TFMT, Cat# HY-156048), and Aminooxyacetic acid hemihydrochloride (OAA, Cat# HY-107994), were purchased from MedChemExpress. Decibatine (Cat# T1508 / 2353-33-5), *N*-(3-aminopropyl)cyclohexylamine (Cat# T35878), and Tazemetostat (Cat# T1788 / 1403254-99-8) were obtained from Targetmol. Buthionine sulfoximine (BSO, Cat# BML-FR117) was purchased from Enzo Life Sciences. The PHGDH inhibitor (BI-4924) was provided by the Boehringer Ingelheim under the OpnMe Initiative. 3-deazaneplanocin A HCl (DzNep, Cat# A13947) was purchased from Adooq Biosciences. 5’-deoxy-5’-methylthioadenosine (MTA, Cat# Ab141257) was obtained from Abcam. Adenine (Cat# 400130.001) was from Pharma Waldhof. Methionine (Cat# 64320) and Cysteine (Cat# 30090) were from Fluka Biochemika. Analytical grade methanol (Cat# 10097370) and acetonitrile (Cat# 12759600) were from J.T. Baker™. FastDigest Esp3I enzyme (Cat# FD0454) was from Thermo Scientific. The wild-type Chikungunya virus (CHIKV^WT^) and CHIKV^Gluc^ were provided as a kind gift from Prof. Juana Díez (Universitat Pompeu Fabra, Barcelona).

### Cell culturing

CHIKV permissive cells: Huh7.5.1 (a kind gift from Prof. Juana Díez (Barcelona)), Vero E6 (ATCC, Cat# ATCC CRL-1586), and HEK293T (a kind gift from Bettina Hinkelmann) cells were maintained in 1× Dulbecco’s modified eagle’s medium (DMEM) (+ GlutaMAX™-I, [+] 4.5g/L D-Glucose and [-] Pyruvate, Gibco™, UK, CAT# 2533742) supplemented with 10% heat-inactivated fetal bovine serum (FBS) and cultured at 37°C and 10% CO_2_. Cells were harvested for experiments on reaching ∼ 90 – 100% confluence.

### Metabolite extractions and profiling

All CHIKV infections were performed following approved protocols in a S3** biosafety level laboratory. Adherent 3 × 10^6^ cells/well in 6-well plates were mock infected or infected with CHIKV at the indicated multiplicity of infection (MOI) for 1 h, and the inoculum subsequently washed off with warm 1× PBS (pH 7.4). Pre-warmed DMEM + 10% FBS (37°C; 1-mL/well) was then added to each well and CHIKV infection allowed for 8 h early time point of viral replication. Intracellular metabolites were extracted as previously described^15^. In brief, spent growth medium was aspirated from each well and cells washed with 1 mL pre-warmed of ammonium carbonate (75 mM; pH 7.2). Metabolism was immediately quenched with 1-mL cold 60% MeOH supplemented with 1× PBS (pH 7.4). After aspiration, metabolites were extracted from each well for 1 h (at –20°C) with cold 400 µL MeOH:ACN:H_2_O (2:2:1 (v/v/v); LC-MS Analytical grade) spiked with internal standards: glipizide (3 µg/mL), naproxen (8 µg/mL), and nortrityline (1 µg/mL). The first batch of metabolites was collected into 1.5-mL Eppendorf tubes, and a second extraction performed, but this time, the cells were scrapped from the well plates and resuspended in the extraction medium for an additional 1 h. After pooling the extracted metabolites, any potential CHIKV contaminants were heat inactivated at 72°C for 30 min^97,98^. Then the samples were centrifuged for 10 min before transferring the supernatants into clean 1.5-mL Eppendorf tubes for SpeedVac concentration. The dried samples were then resuspended in 20 µL 5% acetonitrile/water with 0.1% formic acid solution and transferred into HPLC glass vials for high-resolution LC-MS analyses.

Metabolites were profiled by a high-resolution hydrophilic interaction chromatography mass spectroscopy (HILIC – MS) mode. The liquid chromatography was interfaced with a maXis HD UHR-TOF mass spectrometer (Bruker, Germany) fitted with a Gemini 3 µm NX-C18 column (110Å, 50 × 4,6 mm, Phenomenex) running on a MS^2^ positive mode as previously adapted ^99^. Samples were injected at a flow rate of 300 µl/min and analyzed for 30 min at a gradient elution of solvent A (water in 0.1% formic acid) and solvent B (acetonitrile in 0.1% formic acid). The following gradient conditions were used: 0 – 2 min (1% solvent B), followed by a linear gradient between 2 – 20 min (1% solvent B), 20 – 29 min (100% solvent B) after which the column was washed with 100% solvent B and the starting conditions restored with 1% solvent B. During the analysis, the column temperature was maintained at 40°C and an internal calibration with sodium formate performed for the first 0.3 min of every run. Metabolites data were acquired using electrospray ionization (ESI) under full scan mass range of 50 – 1500 m/z. Collision-induced dissociation was used to fragment the ions. The source parameters were set as follows: capillary voltage of 4500 V, nebulizer pressure at 4.0 bar, dry heater at 200°C, and dry gas at 9.0 l/min. Analyses of the acquired raw data were performed on MetaboScape® 2022 software. For annotations, we considered metabolites between retention times (RTs) 0.3 min ≥ RTs ≤ 25 min filtered applying the parameters outlined in the data analyses section. The metabolites were putatively identified by matching the accurate *m/z*, MS/MS fragmentation patterns, and retention times with those of in-house library (600 metabolites) as chemical standards, Bruker MetaboBase (482025 metabolites), and Bruker HMDB metabolite library (824 metabolites).

### Metabolic stable isotope labeling using L-Met (^13^C-methyl) and L-Cys-^13^C_3_, ^15^N. **3**

×10^6^ Huh7.5.1 cells in 6-well plates were infected with CHIKV^WT^ at MOI 2 for 1 h and the unbound inoculum aspirated. The cells were then washed twice with warm PBS and Met-Cys free DMEM (Gibco™, Cat# 21013024) supplemented with 200 µM of L-Met (^13^C –methyl) (Sigma-Aldrich, Cat# 299146) or L-Cys-^13^C_3_, ^15^N (MedChemExpress, Cat# HY-Y0337S) added. The cells were incubated for 8 h and metabolites extracted for profiling. As control, infected cells were provided with complete medium, non-infected cells with stable isotope fortified media, or complete medium. Metabolites extraction and mass spectrometry profiling were carried out as described above.

#### Analyses of RNA modifications

To quantitate the tRNA modifications by LC-MS-based detection of ribonucleosides, total tRNA was isolated using NucleoSpin miRNA kit (Macherey-Nagel; Cat# 740971). 1 µg of each sample was subjected to nucleoside digestion using nucleoside digestion mix (NEB, #M0649), and analyzed as previously described^100,101^ using Agilent 1290 LC system interfaced with ABSciex Qtrap 6500 mass spectrometer. The samples were separated using LC Chromolith High Resolution RP-18 end capped column (Supelco, 50 × 4.6 mm) under the following parameters; flow rate 1.8 mL/min, injection volume of 5 µL, column temperature of 30°C, and a total run time of 2.5 min. Samples were run on a gradient of solvent A: water (with 0.1% formic acid) and solvent B: acetonitrile (with 0.1% formic acid) conditioned as follows, 0 – 0.5 min (5% solvent B), 1.5 – 1.7 min (100% solvent B), and 1.8 – 2.5 min (1% solvent B). The MS/MS was operated in MRM positive mode under set parameters of curtain gas (20 psi, Nitrogen), ion spray voltage: 5500V, temperature of 400°C, compressed air (Gas 1: 45 psi, Gas 2: 65 psi), and entrance potential of 10V. The mass transitions for mcm^5^U were quantified from 317.06 (+1) to 185 Da (declustering potential: 56V, collision energy: 13V, and cell exit potential: 10V). For mcm^5^s^2^U quantifications, fragmentations were monitored from 333.03 (+1) to 200.9 Da (declustering potential: 1V, collision energy: 13V, cell exit potential: 12V) or 333.03 (+1) to 169.1 Da (at declustering potential: 1V, collision energy: 25V, cell exit potential: 10V). Dwell time for all transitions was set at 35 msec. Standard ribonucleosides were purchased from Biosynth^®^ (mcm^5^U, Cat# NM45525 and mcm^5^s^2^U, Cat# NM159492) processed for LC-MS quantification, and further confirmations made from the modomics database (https://iimcb.genesilico.pl/modomics/modifications). All the MRM measurements were quantified using the freely-accessible Skyline program (MacCoss Laboratory, University of Washington).

For m^6^A quantification, total RNA was extracted from CHIKV-infected or control cells using TRIzol® reagent (Invitrogen, Cat #15596026) following manufacturer’s instructions. Equal RNA amounts (200 ng) were probed for relative m^6^A modifications using EpiQuik^™^ m^6^A RNA methylation Quantification kit (Fluorometric; EpigenTeK, Cat #P-9008) as per manufacturer’s guidelines.

#### Nutrient deprivation and repletion assays

Adherent Huh7.5.1 cells pre-seeded at a density of 1×10^5^ per well for 24 h were infected with CHIKV^WT^ for 1 h at a MOI of 0.05. A subset of the CHIKV^WT^-infected cells were incubated with; (i) methionine-free DMEM + 10% FBS (+ 200 µM cysteine), (ii) cysteine-free DMEM (+ 200 µM methionine and 10% FBS), (iii) methionine-free DMEM (+ 50 µM methionine and 10% FBS), (iv) methionine-free DMEM (+ 100 µM methionine), methionine-free DMEM (+ 200 µM methionine), and methionine-cysteine free DMEM (+ 10% FBS). For all media virus titres were determined by plaque assays (see below) and results were compared to CHIKV virus titres obtained from complete DMEM medium. Time-of-deprivation experiments were performed by variation of the indicated time points of the media change after the start of the infection.

#### Plaque assays

Viral titre quantifications were performed by infecting adherent 1 × 10^5^ Vero E6 cells/well seeded in 48-well plates, with harvested supernatants to a fold-dilution of 10^-5^. Following the aspiration of viral inoculum and subsequent wash-off with 1× phosphate buffer, 200 µL of 1% methylcellulose (Sigma-Aldrich, Cat# M7027) were added to each well and plates were incubated under standard conditions for 72 h. The cells were fixed with 6% paraformaldehyde and stained for 5 min with 0.5% crystal violet in 20% methanol. After rinsing twice with water, the number of plaques were counted and viral titres calculated.

#### CRISPR-Cas9 experiments

Huh7.5.1 cells stably expressing target sgRNAs, or control were generated using a LentiCRISPRv2 construct (purchased from Addgene #52961). Target sgRNA sequences were designed using Custom Alt-R™ CRISPR-Cas9 guide RNA platform (https://eu.idtdna.com/site/order/designtool/index/CRISPR_CUSTOM) and its reverse complement generated on https://www.bioinformatics.org/sms/rev_comp.html (sequences are provided in Table S3). To facilitate cloning onto Esp3I-digested LentiCRISPRv2 plasmid, custom design overhangs; CACCG and AAAC were attached to the 5ʹ-end of sgRNA and the reverse complement sequences, respectively. Annealed oligos were ligated into a linearized LentiCRISPRv2 plasmid using Quick Ligation™ kit (NEB, #M2200S) and co-transfected with psPAX2 (Addgene #12260) and pMD2.G (Addgene, #12259) plasmids into HEK293T cells using X-tremeGene™ 9 DNA-transfection reagent (Roche, Cat# XTG9-RO; 06365779001). 48 h later, the lentiviral supernatants were harvested and filtered with Sartorius Minisart® syringe filter (pore size: 0.45 µm) for transducing Huh7.5.1 cells. In presence of 8 µg/mL polybrene (EMD Millipore Cat# TR-1003-G), the cells were transduced for 48 h following a medium exchange with 5 µg/mL puromycin to allow selection of successful knockout. After another 48 h, the stable cells were expanded for downstream assays following a random confirmation by qPCR. Knockout cells were infected with CHIKV^Gluc^ at MOI of 2 for 48 h, and supernatants harvested for viral quantification.

#### Antiviral assays

Stock solutions of compounds at 10 mM were reconstituted in DMEM + 10% FBS to desired concentrations for assays in 96-well plates. Cells seeded at a density of 2 × 10^4^/ well were allowed for 24 h adherence, after which, the overnight spent medium was aspirated and replaced with 50 µL of fresh DMEM containing respective screen compounds. A reporter CHIKV possessing a Gaussian luciferase (CHIKV^Gluc^) at a MOI 0.01 was added to respective wells alongside 0.1% DMSO non-treated and non-infected controls. Plates were incubated at standard conditions of 37°C and 5% CO_2_ for 72 h after which 8 µL supernatants were aliquoted into sterile 96-well plates containing 10 µL *Renilla* luciferase assay reagent (Promega, Cat# E2810). Luminescence intensities were read out using BioTek Synergy 4.1 microtitre plate reader. Ribavirin served as a positive reference antiviral agent. For cell viability assays, CellTiter-Glo^®^ reagent (Promega, Cat# G7570) was used as per manufacturer’s instructions.

#### Total RNA extraction and qRT-PCR analyses

To analyze cellular gene expression changes under various treatment conditions, total RNA was extracted using TRIzol® reagent (Invitrogen Cat #15596026) following manufacturer’s instructions. The concentrations of the extracted total RNA samples were quantified using a Nanodrop spectrophotometer. cDNA was subsequently synthesized from 1 µg RNA using the Thermo Scientific’s Maxima First Strand synthesis kit for RT-qPCR (Cat #K1641) or RevertAid First Strand synthesis kit (Cat #K1622). All PCR primer sets are provided in Table S4 (designed using the publicly accessible Primer3 webtool; https://primer3.ut.ee/).

The gene expression analyses were processed using 2× iTaq™ Universal SYBR^®^ Green Supermix (Bio-Rad, #1725124) following manufacturer’s guidelines. The data was acquired using QuantStudio™ Design & Analysis Software (v1.5.2) running on a QuantStudio 3 qPCR system. qPCR amplifications were programmed as follows: a hotstart activation step at 95°C for 30 sec, 40 cycles of denaturation step at 95°C for 15 sec, annealing step at 60°C for 30 sec, and then finally by a melt curve step 65°C to 95°C at system default settings. Each gene was analyzed in triplicate and human GAPDH (5′-3′ Fwd: GTTCGACAGTCAGCCGCAT, Rev: GGAATTTGCCATGGGTGGA) served as the endogenous normalization control. Relative fold changes were analyzed using the 2^^-ΔΔCT^ method^102^.

To quantify the CHIKV genome copies, total intracellular RNA was isolated and processed as described above. Extracellular viral RNA was purified from supernatants using PureLink™ Viral RNA/DNA mini kit (Invitrogen™, #12280050) as per the manufacturer’s guidelines. cDNA samples, which resulted from reverse transcription from 1 µg of each purified RNA sample were used as templates for amplification of CHIKV E1. CHIKV E1 primer pair used (5ʹ-3ʹ): Fwd: ACGCAGTTGAGCGAAGCAC, Rev: CTGAAGACATTGGCCCCAC. Absolute viral genomic copies were derived from a corresponding amount on a standard curve generated using a synthetic CHIKV E1 gBlock gene fragment (Integrated DNA Technologies).

#### Cellular ROS determination

CHIKV-infected cells treated with 10 µM DzNep, 10 µM Adox, Met-Cys free, or an infection control were processed for an intracellular ROS assay using 2’,7’-dichlorodihydrofluorescein diacetate (H_2_DCFDA) reagent (Invitrogen™, Cat # D399) according to the manufacturer’s guidelines.

#### Indirect immunofluorescence microscopy

Cells seeded on coverslips were infected with CHIKV and treated with DzNep, Adox, Met-Cys free medium, or DMSO for 8 hpi. The culture medium was aspirated from each well and cells fixed with 4% formaldehyde for 10 min. Following washing off of formaldehyde with 1× PBS, the cells were permeabilized for 15 min with 0.1% Triton X-100 (in 1× PBS) and blocked with 2% BSA added for 1 h. The cells were subsequently incubated overnight at 4°C with CHIKV nsP2 antibody (Thermo Fisher #PA5-143493; dilution 1:500). Following discard of the antibody and rinsing three times with 1× PBS, a fluorescent labeled secondary antibody; Alexa Fluor 488 goat anti-rabbit IgG H&L (Abcam, Cat# Ab150077, dilution 1:500) was added to each coverslip and incubated in dark at room temperature for 1 h. The stained cells were finally washed three times with 1× PBS and mounted onto glass slides with ProLong™ Gold Antifade mountant with DAPI (Molecular probes, #P36941). Images were captured using EVOS™ M5000 imaging system.

#### Statistical data analysis

Analyses were performed on Graphpad prism 9.3.1 and RStudio (version 2023.03.0) running on R (version 4.2.3) softwares. The *Student’s t*-tests were applied for comparison of two independent treatment groups. Dose-response curves for estimating inhibitory concentrations were generated through a 4-parameter variable slope regression fit modelling. Source metabolomics data were analyzed on MetaboScape® 2022 software. The data were processed by applying MCube T-Rex 3D Processing Workflow method (version 1.91). The following filter parameters were applied: minimum # features for extraction – 25%, presence of features in minimum # of analyses – 60%, and 50% to 83% presence of features in a sample group. For all the experimental analyses, *p* values of less than 5% were considered statistically significant. The asterisks designate *p* ≤ 0.05 (*), *p* ≤ 0.01 (**), *p* ≤ 0.001 (***), *p* ≤ 0.0001 (****), and *p* > 0.05 (ns – not significant). Unless mentioned, all the data are presented as mean ± standard deviation (s.d.) of experimental replicates.

#### Data availability

All data to support the conclusions are provided in the manuscript and its supplementary information, and available upon formal request to Ursula Bilitewski.

## Acknowledgements

This study was funded by the Alexander von Humboldt-Stiftung awarded to J.M.M. under the Alexander von Humboldt – Bayer Science Foundation fellowship program. The funding body had no influence over the study design, data collection and analyses, decision to publish, and/or preparation of the manuscript. Assistance of Dr. Kai Schulze in S3** laboratory biosafety guidance and protocol approvals, and excellent facilitation of Britta Kohler and Josefin Koch is hereby acknowledged. We extend our sincere gratitude to Claudia Soltendieck, Dr. Aditya Shekhar, and Konstanze Luckhardt for assistance with ordering of experimental supplies.

## Author contributions

J.M.M. and U.B.* conceived and designed the study. J.M.M performed the experiments, analyzed data, and wrote the initial manuscript draft. M.F. facilitated cell culture maintenance and consumables ordering. H.O.-M., U.B., and R.F. assisted in metabolomics data acquisitions. P.S., M.B., and U.B.* provided supervision, data curation, editing, and infrastructural resource facilitation. All authors revised and agreed on the final version.

## Supplemental information titles and legends

**Fig. S1:**
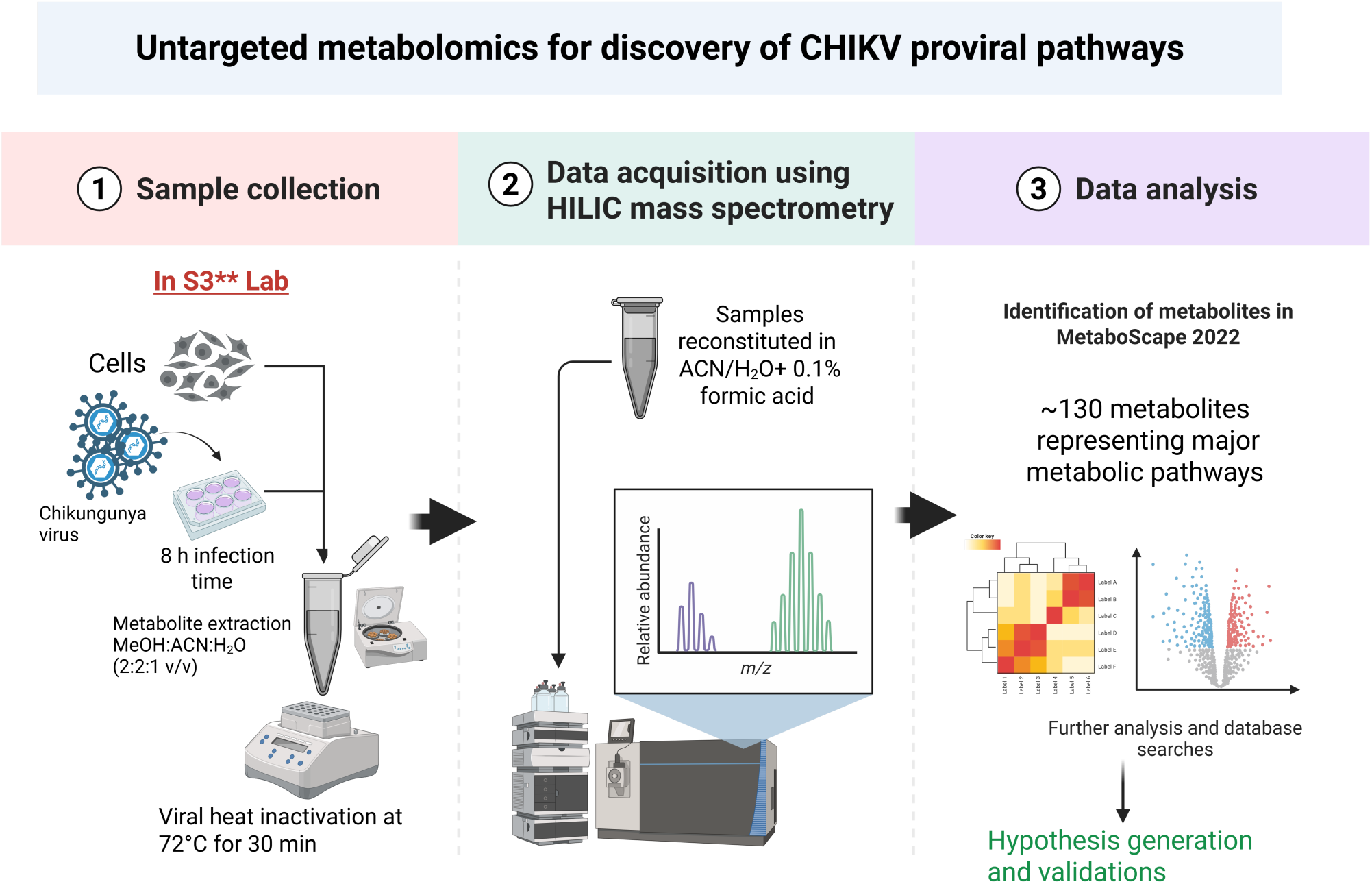
Metabolomics workflow. CHIKV-infected cells were subjected to metabolite extraction 8 hours post infection (hpi) using MeOH:ACN:H_2_O (2:2:1). The extracts were heat-inactivated for 30 min at 72°C, centrifuged, concentrated in a speedvac, and reconstituted for LC-MS profiling. Mass spectra were analyzed using MetaboScape software to link features to metabolites.

**Fig. S2:**
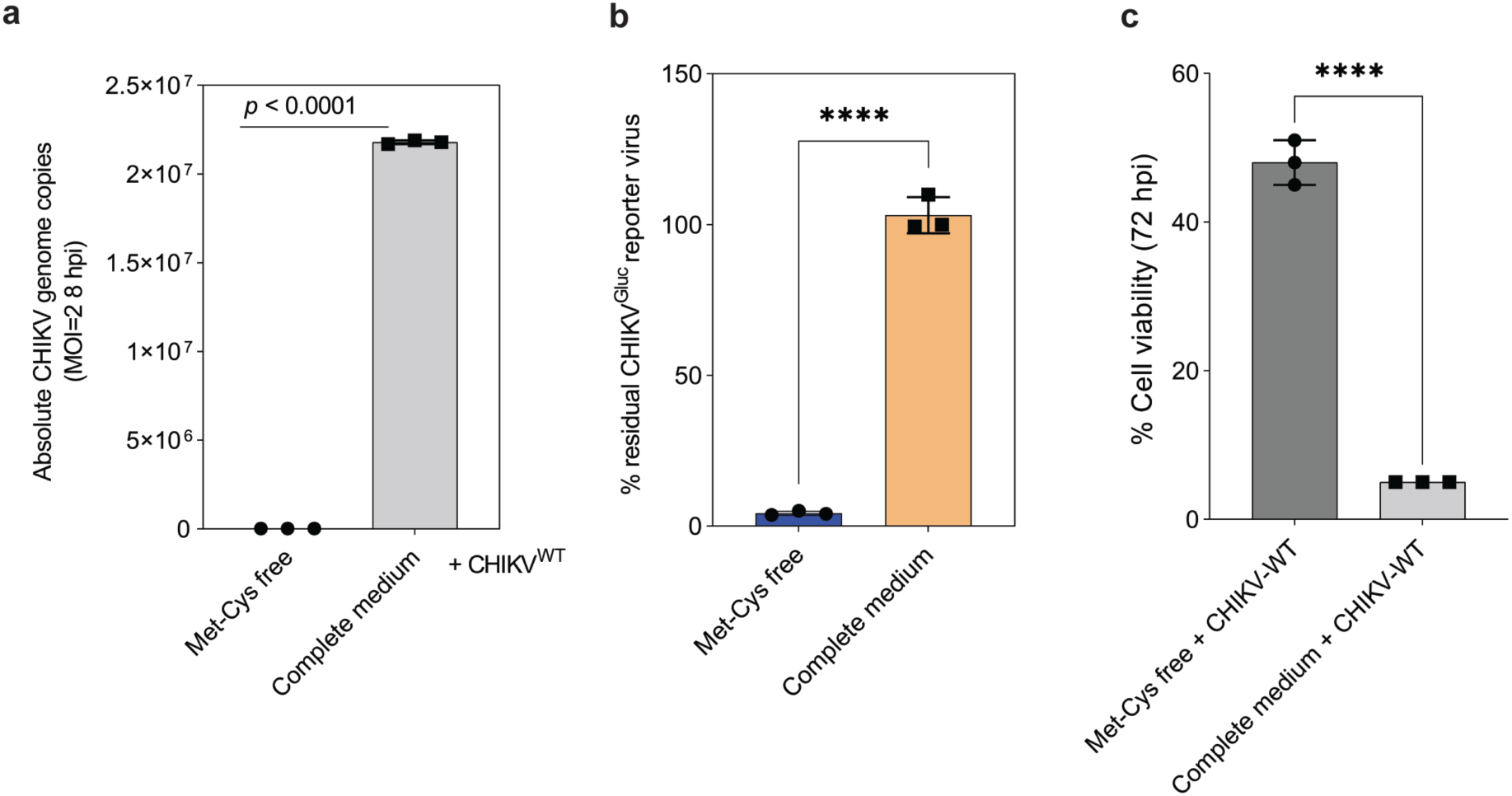
Met-Cys deprivation effects on CHIKV replication at different time points. Absolute CHIKV genome copies quantification by qPCR (8 hpi), reporter virus luciferase (72 hpi), and cell viability-based cytopathic effects (72 hpi). *Student*’s *t*-test, **a**, t = 377, df = 2, *p* < 0.0001, **b**, t = 28.47, df = 4, *p* < 0.0001, **c**, t = 24.83, df = 4, *p* < 0.0001.

**Fig. S3:**
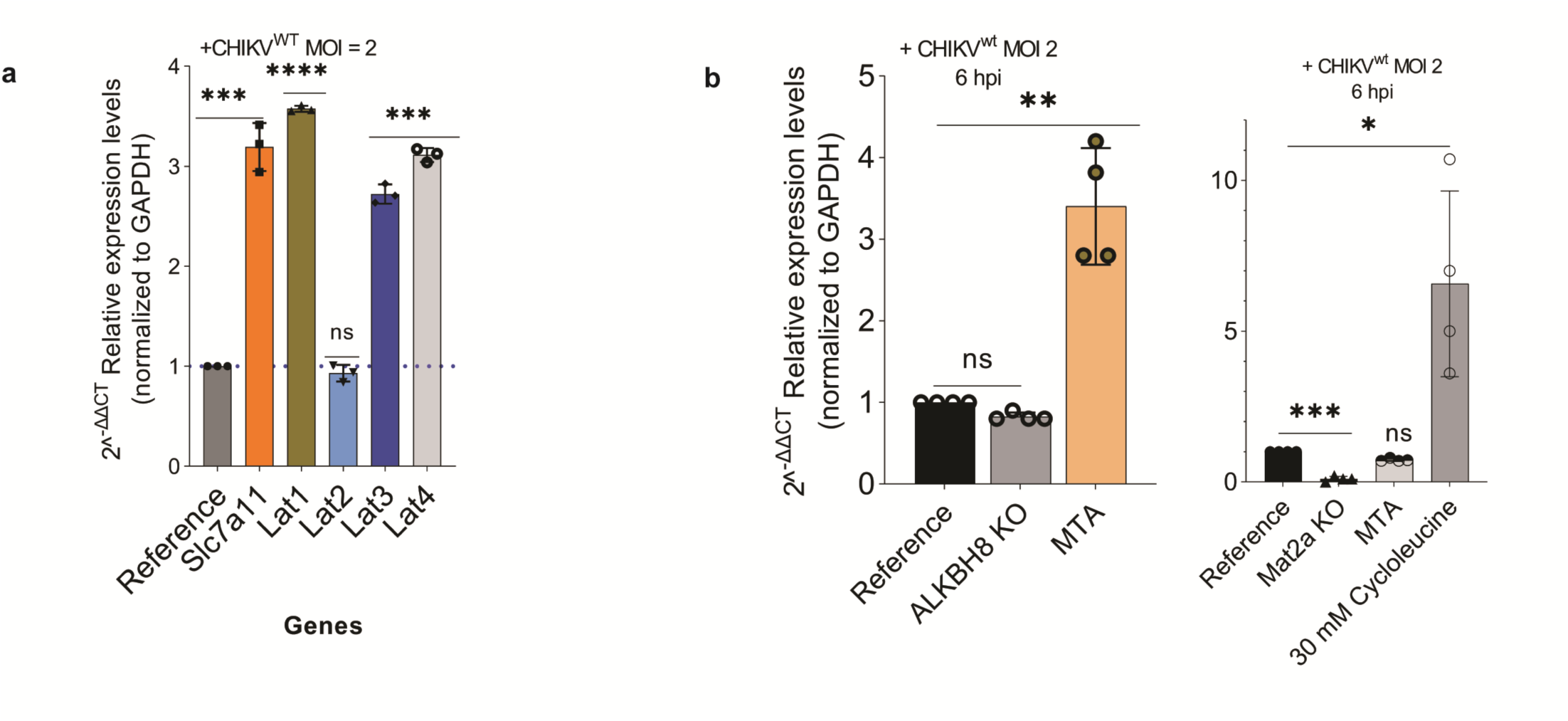
Fold changes in expression of solute carriers for Met and Cys under Met-Cys deprivation. (**a**) and Mat2a-MTA-ALKBH8 relationship analysis under CHIKV^WT^ MOI of 2 infection challenge (**b**). Data presented as mean ± s.d. of three (**a**) and four (**b**) experimental replications. *Student’s t*-test, **a**, *Slc7a11* vs reference, t = 15.9, df = 2, *p* = 0.0039, *Lat1*, t = 146.5, df = 2, *p* < 0.0001, *Lat2*, t = 1.458, df = 2, *p* = 0.2823, *Lat3*, t = 31.25, df = 2, *p* = 0.001, *Lat4*, t = 52.99, df = 2, *p* = 0.0004. **b,** *ALKBH8*: MTA vs reference, t = 11.34, df = 6, *p* < 0.0001, Mat2a KO vs reference, t = 22.05, df = 6, *p* < 0.0001. *Mat2a*: MTA vs reference, t = 6.722, df = 3, *p* = 0.0067. ns – not significant.

**Fig. S4:**
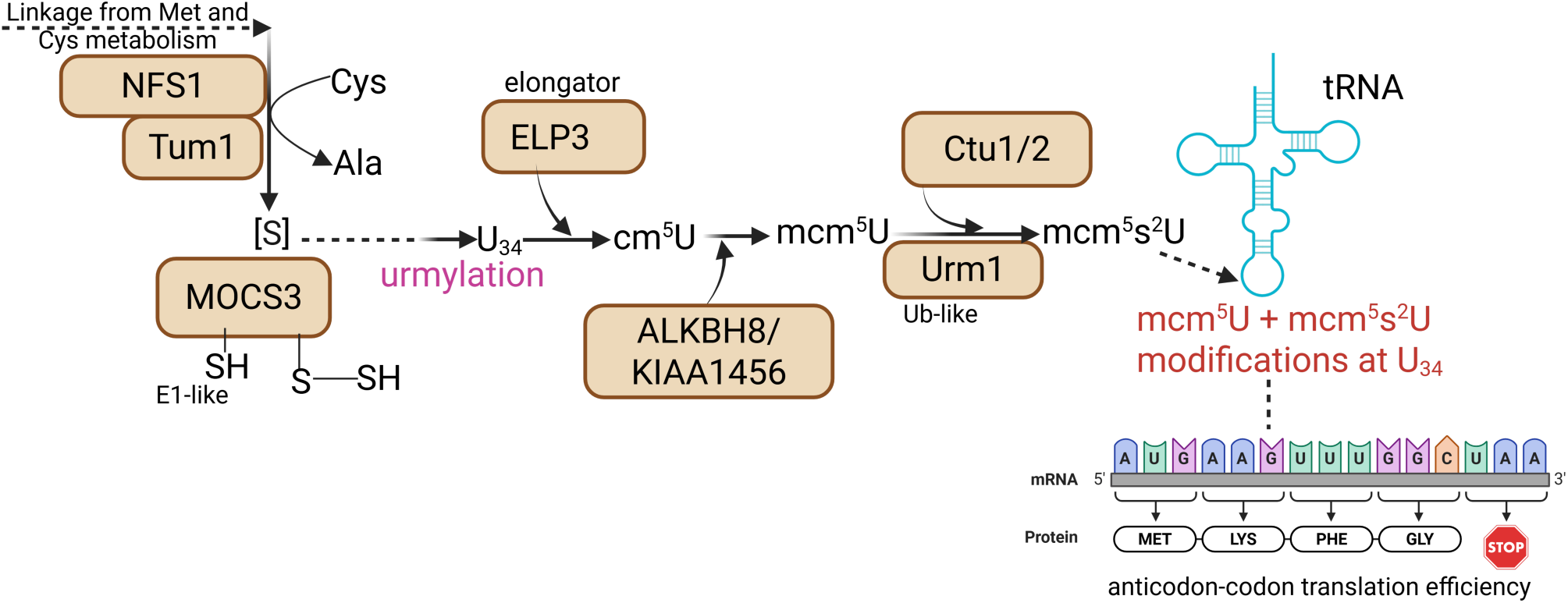
Sulfur relay system in eukaryotic cells. Sulfur from methionine and cysteine pathways is converted into persulfide through desulfuration activities of Nfs1 and TUM1. MOCS3 and Urm1 acts as the sulfur carriers, which through urmylation process facilitates adenylation and thiocarboxylation to give URM1-COSH (activated sulfur donor). This sulfur is then transferred to uridine at wobble site 34 of tRNA (U_34_) through three chemical reactions: formation of 5-carbamoylmethyluridine (cm^5^U) catalyzed by elongator complex protein 3 (ELP3), followed by methylation of cm^5^U into 5-methoxycarbonylmethyluridine (mcm^5^U) by ALKBH8/KIAA1456, and finally 2-thiolation of mcm^5^U into mcm^5^s^2^U through actions of Urm1 and cytosolic thiouridylases Ctu1/2. ALKBH8, Alkylated DNA repair protein alkB homolog 8; Ctu1/2, Cytoplasmic tRNA 2-thiolation protein 1/2; ELP3, Elongator complex protein 3; KIAA1456, Probable tRNA methyltransferase 9B (trmt9b); MOCS3, Molybdenum cofactor synthesis protein 3; NFS1, Cysteine desulfurase; TUM1, Thiosulfate sulfurtransferase 1; Urm1, Ubiquitin-related modifier 1. Re-designed from KEGG database on Biorender.com

**Fig. S5:**
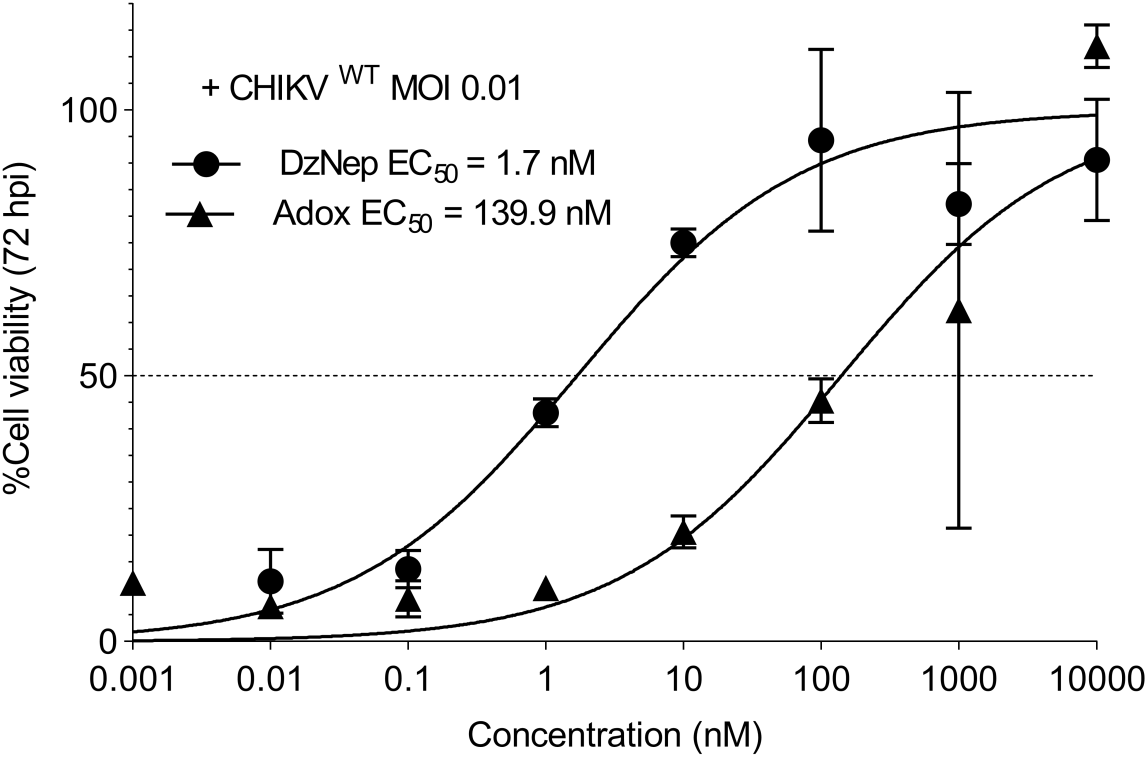
Confirmation of antiviral activity of DzNep and Adox against wild-type CHIKV. Cell viability was performed after 72 hpi using Cell-Titer Glo® assay.

**Table S1:** Summarized metabolomics data obtained from Huh7.5.1 and Vero E6 cells infected with CHIKV at MOI of 0.2, 2, and 20 for 8 hpi.

**Table S2:**
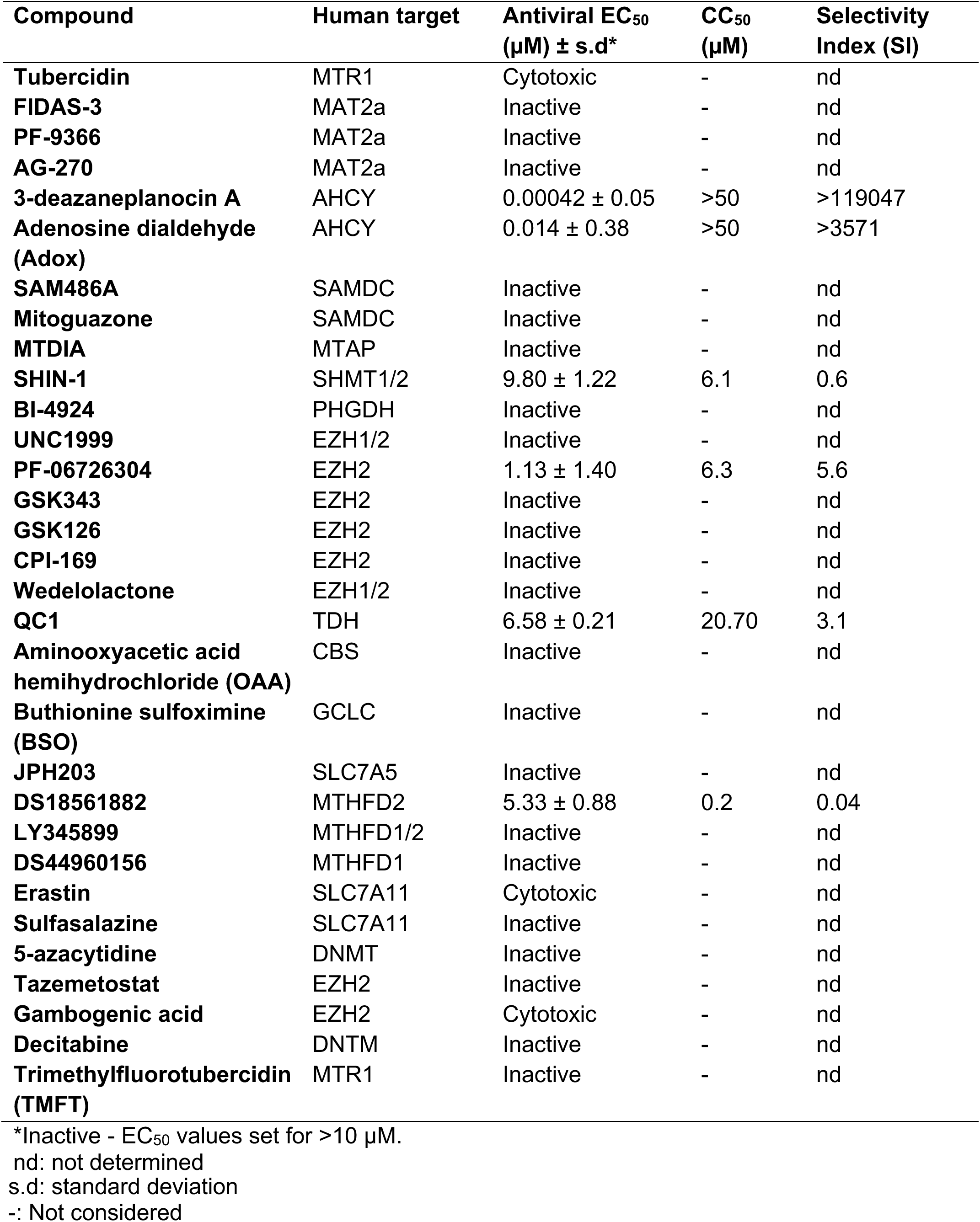
List of compounds tested for antiviral activity against CHIKV and their activity profiles. Abbreviations: AHCY (SAH hydrolase), CBS (Cystathionine β synthase), DNMT (DNA methyltransferase), EZH1/2 (Enhancer of Zeste 1/2), GCLC (glutamate-cysteine ligase), MAT2a (Methionine adenosyltransferase 2A), MTAP (Methylthioadenosine Phosphorylase), MTHFD1/2 (Methylenetetrahydrofolate dehydrogenase 1/2), MTR1 (Methionine synthase), PHGDH (Phosphoglycerate dehydrogenase), SAMDC (*S*-adenosylmethionine decarboxylase), TDH (Threonine dehydrogenase), SHMT1/2 (Serine hydroxymethyltransferase 1/2), SLC7A5 (Solute carrier family 7 member 5), SLC7A11 (Solute carrier family 7 member 11).

**Table S3:**
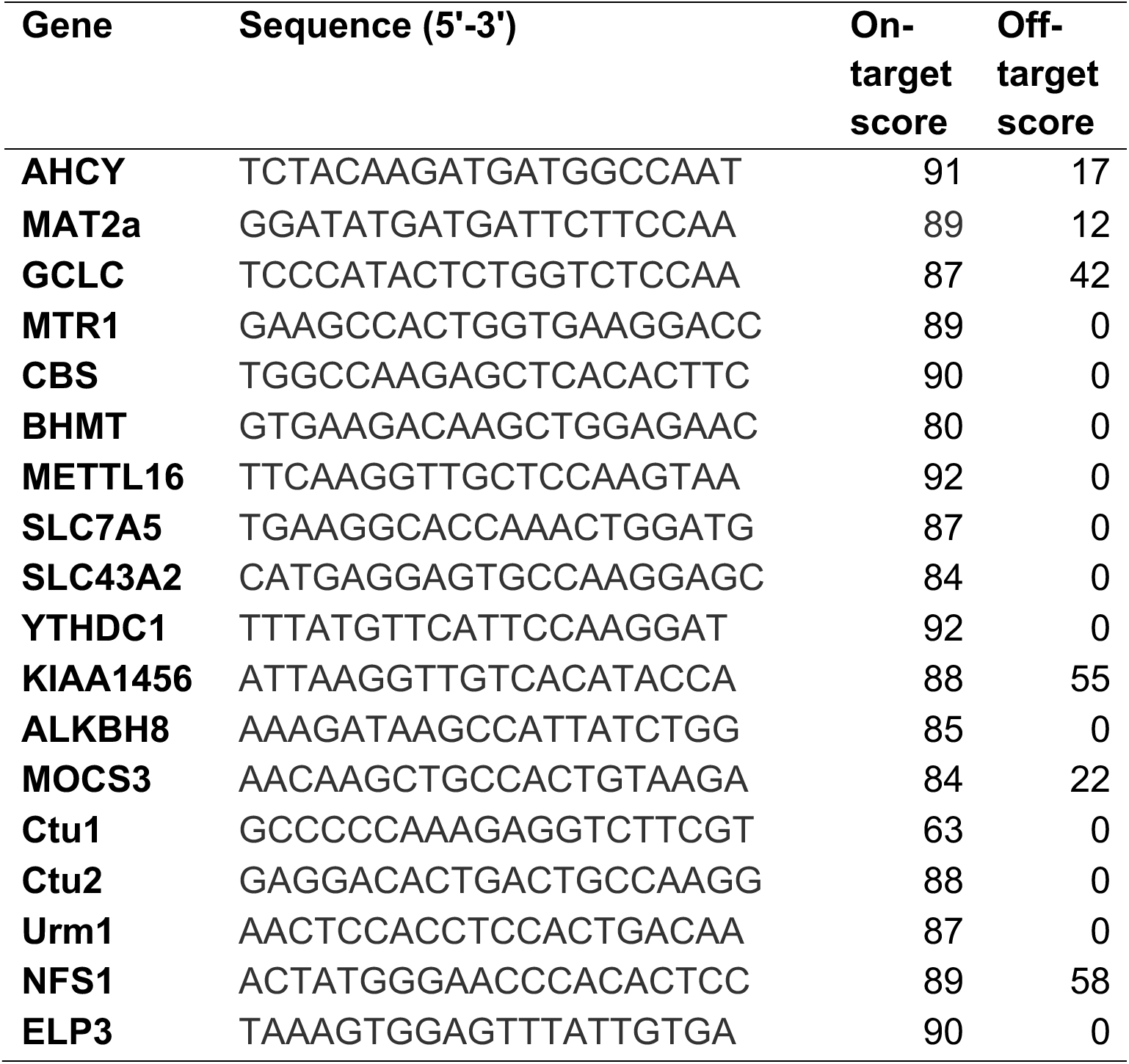
CRISPR sgRNA oligonucleotides. Coding sequences were retrieved from UniProt database. Corresponding sgRNAs targeting the genes were designed on IDT custom-design tool (https://eu.idtdna.com/site/order/designtool/index/CRISPR_CUSTOM).

**Table S4:**
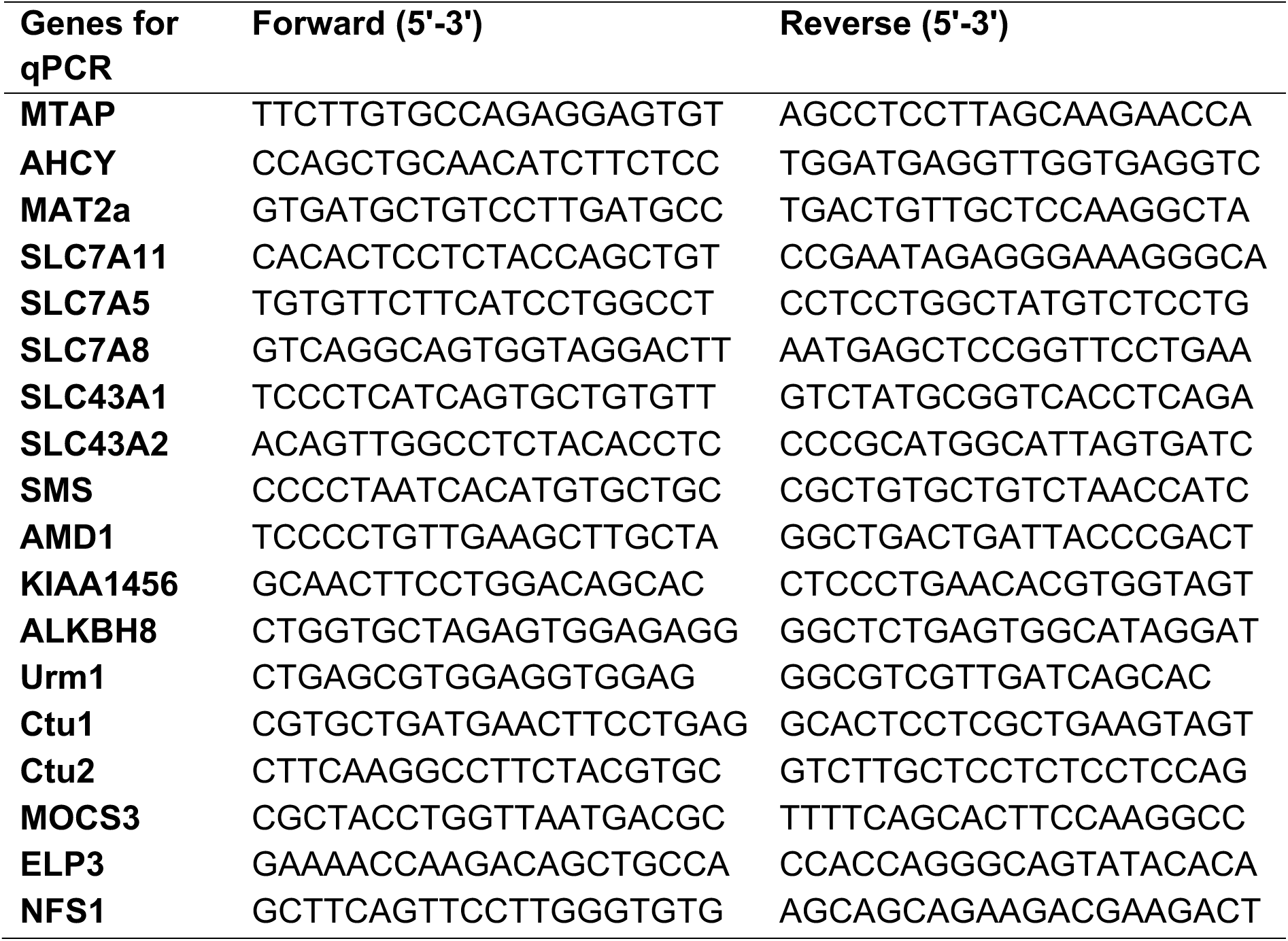
Primer pairs used for qPCR analyses. Sequences of target proteins were retrieved from Uniprot database (https://www.uniprot.org/uniprotkb). The primer pairs were designed on Primer3 input https://primer3.ut.ee/cgi-bin/primer3/primer3web_results.cgi

## Supplementary data files

**Supplementary data file 1 – 3**: Metabolomics dataset of CHIKV-infected cells at MOI of 0.2, 2, and 20, respectively, for Huh7.5.1 and Vero E6 cells.

**Supplementary data file 4**: Stable isotope labelling experiment results.

## References

1. Lumsden, W. An epidemic of virus disease in Southern Province, Tanganyika territory, in 1952–1953 II. General description and epidemiology. Trans R Soc Trop Med Hyg 49, 33–57 (1955).

2. de Souza, W. M. et al. Pathophysiology of chikungunya virus infection associated with fatal outcomes. Cell Host Microbe 32, 606–622.e8 (2024).

3. Pérez-Pérez, M.-J., Delang, L., Ng, L. F. P. & Priego, E.-M. Chikungunya virus drug discovery: still a long way to go? Expert Opin. Drug Discov. 14, 855–866 (2019).

4. Rochlin, I., Ninivaggi, D. V, Hutchinson, M. L. & Farajollahi, A. Climate Change and Range Expansion of the Asian Tiger Mosquito (Aedes albopictus) in Northeastern USA: Implications for Public Health Practitioners. PLoS One 8, e60874 (2013).

5. Liu-Helmersson, J., Rocklöv, J., Sewe, M. & Brännström, Å. Climate change may enable Aedes aegypti infestation in major European cities by 2100. Environ. Res. 172, 693–699 (2019).

6. Kraemer, M. U. G. et al. Past and future spread of the arbovirus vectors Aedes aegypti and Aedes albopictus. Nat. Microbiol. 4, 854–863 (2019).

7. WHO. Global arbovirus initiative: preparing for the next pandemic by tackling mosquito-borne viruses with epidemic and pandemic potential. (2024).

8. Thaker, S. K., Ch’ng, J. & Christofk, H. R. Viral hijacking of cellular metabolism. BMC Biol. 17, 59 (2019).

9. Goodwin, C. M., Xu, S. & Munger, J. Stealing the Keys to the Kitchen: Viral Manipulation of the Host Cell Metabolic Network. Trends Microbiol. 23, 789–798 (2015).

10. Allen, C. N. S., Arjona, S. P., Santerre, M. & Sawaya, B. E. Hallmarks of Metabolic Reprogramming and Their Role in Viral Pathogenesis. Viruses vol. 14 602 at 10.3390/v14030602 (2022).

11. Thaker, S. K. et al. Differential Metabolic Reprogramming by Zika Virus Promotes Cell Death in Human versus Mosquito Cells. Cell Metab. 29, 1206–1216.e4 (2019).

12. Denolly, S. et al. Zika virus remodelled ER membranes contain proviral factors involved in redox and methylation pathways. Nat. Commun. 14, 8045 (2023).

13. Mullen, P. J. et al. SARS-CoV-2 infection rewires host cell metabolism and is potentially susceptible to mTORC1 inhibition. Nat. Commun. 12, 1876 (2021).

14. Zhang, Y. et al. SARS-CoV-2 hijacks folate and one-carbon metabolism for viral replication. Nat. Commun. 12, 1676 (2021).

15. Gassen, N. C. et al. SARS-CoV-2-mediated dysregulation of metabolism and autophagy uncovers host-targeting antivirals. Nat. Commun. 12, 3818 (2021).

16. Fontaine, K. A., Sanchez, E. L., Camarda, R. & Lagunoff, M. Dengue virus induces and requires glycolysis for optimal replication. J. Virol. 89, 2358–2366 (2015).

17. Chen, Q. et al. Metabolic reprogramming by Zika virus provokes inflammation in human placenta. Nat. Commun. 11, 2967 (2020).

18. Pang, H. et al. Aberrant NAD+ metabolism underlies Zika virus–induced microcephaly. Nat. Metab. 3, 1109–1124 (2021).

19. Yau, C. et al. Dysregulated metabolism underpins Zika-virus-infection-associated impairment in fetal development. Cell Rep. 37, 110118 (2021).

20. Heaton, N. S. & Randall, G. Dengue Virus-Induced Autophagy Regulates Lipid Metabolism. Cell Host Microbe 8, 422–432 (2010).

21. Dias, S. S. G. et al. Metabolic reprogramming and lipid droplets are involved in Zika virus replication in neural cells. J. Neuroinflammation 20, 61 (2023).

22. Al-Shalan, H. A. M. et al. Systemic perturbations in amino acids/amino acid derivatives and tryptophan pathway metabolites associated with murine influenza A virus infection. Virol. J. 20, 270 (2023).

23. Greene, K. S. et al. Glutamine metabolism is essential for coronavirus replication in host cells and in mice. J. Biol. Chem. 301, 108063 (2025).

24. Mitra, S. et al. Rotavirus rewires host cell metabolic pathways toward glutamine catabolism for effective virus infection. Gut Microbes 16, 2428425 (2024).

25. Pant, A. & Yang, Z. Asparagine: An Achilles Heel of Virus Replication? ACS Infect. Dis. 6, 2301–2303 (2020).

26. Huang, R., et al. Borna disease virus infection perturbs energy metabolites and amino acids in cultured human oligodendroglia cells. (2012).

27. Shrinet, J., Shastri, J. S., Gaind, R., Bhavesh, N. S. & Sunil, S. Serum metabolomics analysis of patients with chikungunya and dengue mono/co-infections reveals distinct metabolite signatures in the three disease conditions. Sci. Rep. 6, 36833 (2016).

28. Lee, C.-D. & Tu, B. P. Metabolic influences on RNA biology and translation. Crit. Rev. Biochem. Mol. Biol. 52, 176–184 (2017).

29. Shu, X. E., Swanda, R. V & Qian, S.-B. Nutrient control of mRNA translation. Annu. Rev. Nutr. 40, 51–75 (2020).

30. Honarmand Ebrahimi, K., et al. Iron–sulfur clusters as inhibitors and catalysts of viral replication. Nat. Chem. 14, 253–266 (2022).

31. Si-Ying, L. A., Rebeca, B.-K. & J., W. S. P. A Genome-Wide Small Interfering RNA Screen Identifies Host Factors Required for Vesicular Stomatitis Virus Infection. J. Virol. 88 (*15*), 8355–8360 (2014).

32. Panda, D. et al. RNAi screening reveals requirement for host cell secretory pathway in infection by diverse families of negative-strand RNA viruses. Proc. Natl. Acad. Sci. 108, 19036–19041 (2011).

33. Bergant, V. et al. Attenuation of SARS-CoV-2 replication and associated inflammation by concomitant targeting of viral and host cap 2’-O-ribose methyltransferases. EMBO J. 41, e111608 (2022).

34. Jungfleisch, J. et al. CHIKV infection reprograms codon optimality to favor viral RNA translation by altering the tRNA epitranscriptome. Nat. Commun. 13, 4725 (2022).

35. Karlas, A. et al. A human genome-wide loss-of-function screen identifies effective chikungunya antiviral drugs. Nat. Commun. 7, 11320 (2016).

36. Kim, B. et al. Discovery of Widespread Host Protein Interactions with the Pre-replicated Genome of CHIKV Using VIR-CLASP. Mol. Cell 78, 624–640.e7 (2020).

37. Reis, Erik V. S., Damas, Beatriz M., Mendonça, Diogo C., Abrahão, Jônatas S., Bonjardim, C. A. In-depth characterization of the Chikungunya virus replication cycle. J. Virol. 96, e01732–21 (2022).

38. Avila, M. A., Garcı’a-Trevijano, E. R., Lu, S. C., Corrales, F. J. & Mato, J. M. Methylthioadenosine. Int. J. Biochem. Cell Biol. 36, 2125–2130 (2004).

39. Ning, T. et al. Newcastle Disease Virus Manipulates Mitochondrial MTHFD2-Mediated Nucleotide Metabolism for Virus Replication. J. Virol. 97, e00016–23 (2023).

40. Roe, B., Kensicki, E., Mohney, R. & Hall, W. W. Metabolomic Profile of Hepatitis C Virus-Infected Hepatocytes. PLoS One 6, e23641 (2011).

41. Mounce, B. C. et al. Interferon-Induced Spermidine-Spermine Acetyltransferase and Polyamine Depletion Restrict Zika and Chikungunya Viruses. Cell Host Microbe 20, 167–177 (2016).

42. Mounce, B. C. et al. Inhibition of Polyamine Biosynthesis Is a Broad-Spectrum Strategy against RNA Viruses. J. Virol. 90, 9683–9692 (2016).

43. Greco, A. et al. S-adenosyl methionine decarboxylase activity is required for the outcome of herpes simplex virus type 1 infection and represents a new potential therapeutic target. FASEB J. 19, 1128–1130 (2005).

44. Schäfer, B. et al. Inhibition of Multidrug-Resistant HIV-1 by Interference with Cellular S-adenosylmethionine Decarboxylase Activity. J. Infect. Dis. 194, 740– 750 (2006).

45. Shirahata, A., Takahashi, N., Beppu, T., Hosoda, H. & Samejima, K. Effects of inhibitors of spermidine synthase and spermine synthase on polyamine synthesis in rat tissues. Biochem. Pharmacol. 45, 1897–1903 (1993).

46. Regenass, U. et al. CGP 48664, a new S-adenosylmethionine decarboxylase inhibitor with broad spectrum antiproliferative and antitumor activity. Cancer Res. 54, 3210–3217 (1994).

47. Martínez-Chantar, M. L. et al. L-Methionine Availability Regulates Expression of the Methionine Adenosyltransferase 2A Gene in Human Hepatocarcinoma Cells: ROLE OF S-ADENOSYLMETHIONINE*. J. Biol. Chem. 278, 19885–19890 (2003).

48. Albers, E. Metabolic characteristics and importance of the universal methionine salvage pathway recycling methionine from 5′-methylthioadenosine. IUBMB Life 61, 1132–1142 (2009).

49. Pendleton, K. E. et al. The U6 snRNA m6A Methyltransferase METTL16 Regulates SAM Synthetase Intron Retention. Cell 169, 824–835.e14 (2017).

50. Shima, H. et al. S-Adenosylmethionine Synthesis Is Regulated by Selective N6-Adenosine Methylation and mRNA Degradation Involving METTL16 and YTHDC1. Cell Rep. 21, 3354–3363 (2017).

51. Navik, U. et al. Methionine as a double-edged sword in health and disease: Current perspective and future challenges. Ageing Res. Rev. 72, 101500 (2021).

52. Bröer, S. & Bröer, A. Amino acid homeostasis and signalling in mammalian cells and organisms. Biochem. J. 474, 1935–1963 (2017).

53. Laxman, S. et al. Sulfur Amino Acids Regulate Translational Capacity and Metabolic Homeostasis through Modulation of tRNA Thiolation. Cell 154, 416–429 (2013).

54. Kalhor, H. R. & Clarke, S. Novel Methyltransferase for Modified Uridine Residues at the Wobble Position of tRNA. Mol. Cell. Biol. 23, 9283–9292 (2003).

55. Songe-Møller, L. et al. Mammalian ALKBH8 Possesses tRNA Methyltransferase Activity Required for the Biogenesis of Multiple Wobble Uridine Modifications Implicated in Translational Decoding. Mol. Cell. Biol. 30, 1814–1827 (2010).

56. Deng, W. et al. Trm9-Catalyzed tRNA Modifications Regulate Global Protein Expression by Codon-Biased Translation. PLOS Genet. 11, e1005706 (2015).

57. Cavallin, I. et al. HITS-CLIP analysis of human ALKBH8 reveals interactions with fully processed substrate tRNAs and with specific noncoding RNAs. RNA 28, 1568–1581 (2022).

58. Gokhale, N. S. et al. N6-Methyladenosine in Flaviviridae Viral RNA Genomes Regulates Infection. Cell Host Microbe 20, 654–665 (2016).

59. Wendt, L. et al. N6-methyladenosine is required for efficient RNA synthesis of Ebola virus and other haemorrhagic fever viruses. Emerg. Microbes Infect. 12, 2223732 (2023).

60. Mertens, P. P. C. & Payne, C. C. The effects of S-Adenosyl methionine (AdoMet) and its analogues on the control of transcription and translation in vitro of the mRNA products of two cytoplasmic polyhedrosis viruses. Virology 131, 18–29 (1983).

61. Furuichi, Y. Allosteric stimulatory effect of S-adenosylmethionine on the RNA polymerase in cytoplasmic polyhedrosis virus. A model for the positive control of eukaryotic transcription. J. Biol. Chem. 256, 483–493 (1981).

62. Wolfe, M. S. & Borchardt, R. T. S-Adenosyl-L-homocysteine hydrolase as a target for antiviral chemotherapy. J. Med. Chem. 34, 1521–1530 (1991).

63. Marquez, V. E. 3-Deazaneplanocin A (DZNep): A drug that deserves a second look. J. Med. Chem. 67, 17964–17979 (2024).

64. Mudgal, R., Mahajan, S. & Tomar, S. Inhibition of Chikungunya virus by an adenosine analog targeting the SAM-dependent nsP1 methyltransferase. FEBS Lett. 594, 678–694 (2020).

65. Miranda, T. B. et al. DZNep is a global histone methylation inhibitor that reactivates developmental genes not silenced by DNA methylation. Mol. Cancer Ther. 8, 1579–1588 (2009).

66. Joubert, P.-E. et al. Chikungunya virus–induced autophagy delays caspase-dependent cell death. J. Exp. Med. 209, 1029–1047 (2012).

67. Camini, F. C. et al. Oxidative stress in Mayaro virus infection. Virus Res. 236, 1–8 (2017).

68. Guo, R. et al. Methionine metabolism controls the B cell EBV epigenome and viral latency. Cell Metab. 34, 1280–1297.e9 (2022).

69. Wang, L. W. et al. Epstein-Barr-Virus-Induced One-Carbon Metabolism Drives B Cell Transformation. Cell Metab. 30, 539–555.e11 (2019).

70. Mounce BC, Olsen ME, Vignuzzi M, C. J. Polyamines and Their Role in Virus Infection. Microbiol. Mol. Biol. Rev. 81, 10.1128/mmbr.00029–17 (2017).

71. Olsen ME, Filone CMRozelle DMire CE, Agans KN, Hensley L, C. J. Polyamines and Hypusination Are Required for Ebolavirus Gene Expression and Replication. MBio 7, 10.1128/mbio.00882–16 (2016).

72. Firpo, M. R. et al. Targeting Polyamines Inhibits Coronavirus Infection by Reducing Cellular Attachment and Entry. ACS Infect. Dis. 7, 1423–1432 (2021).

73. Bigaud, E. & Corrales, F. J. Methylthioadenosine (MTA) Regulates Liver Cells Proteome and Methylproteome: Implications in Liver Biology and Disease*. Mol. Cell. Proteomics 15, 1498–1510 (2016).

74. Yang, X. et al. MAT2A-mediated S-adenosylmethionine level in CD4+ T cells regulates HIV-1 latent infection. Front. Immunol. 12, 745784 (2021).

75. Ebrahimi, H. K. et al. Iron–sulfur clusters as inhibitors and catalysts of viral replication. Nat. Chem. 14, 253–266 (2022).

76. Nunes, A. et al. Emerging Roles of tRNAs in RNA Virus Infections. Trends Biochem. Sci. 45, 794–805 (2020).

77. Maynard, N. D., Macklin, D. N., Kirkegaard, K. & Covert, M. W. Competing pathways control host resistance to virus via tRNA modification and programmed ribosomal frameshifting. Mol. Syst. Biol. 8, 567 (2012).

78. Baquero-Pérez, B. et al. N6-methyladenosine modification is not a general trait of viral RNA genomes. Nat. Commun. 15, 1964 (2024).

79. Lotz, T. S. & Suess, B. Small-molecule-binding riboswitches. Microbiol. Spectr. 6, 10–1128 (2018).

80. DebRoy, S. et al. A riboswitch-containing sRNA controls gene expression by sequestration of a response regulator. Science (80-.). 345, 937–940 (2014).

81. Breaker, R. R. Riboswitches and the RNA world. Cold Spring Harb. Perspect. Biol. 4, a003566 (2012).

82. Marcel, O., Hendrik, H., Rodney, R., Chen, L. & Ben, B. A Riboswitch Regulates RNA Dimerization and Packaging in Human Immunodeficiency Virus Type 1 Virions. J. Virol. 78, 10814–10819 (2004).

83. Dibrov, S. M. et al. Structure of a hepatitis C virus RNA domain in complex with a translation inhibitor reveals a binding mode reminiscent of riboswitches. Proc. Natl. Acad. Sci. 109, 5223–5228 (2012).

84. Bell, C. L. et al. Control of alphavirus-based gene expression using engineered riboswitches. Virology 483, 302–311 (2015).

85. Wang, S. & White, K. A. Riboswitching on RNA virus replication. Proc. Natl. Acad. Sci. 104, 10406–10411 (2007).

86. Ketzer, P., et al. Artificial riboswitches for gene expression and replication control of DNA and RNA viruses. Proc. Natl. Acad. Sci. 111, E554–E562 (2014).

87. Kendall, C. et al. Structural and phenotypic analysis of Chikungunya virus RNA replication elements. Nucleic Acids Res. 47, 9296–9312 (2019).

88. Sherwood, A. V et al. Hepatitis C virus RNA is 5′-capped with flavin adenine dinucleotide. Nature 619, 811–818 (2023).

89. DeVito, S. R., Ortiz-Riaño, E., Martínez-Sobrido, L. & Munger, J. Cytomegalovirus-mediated activation of pyrimidine biosynthesis drives UDP– sugar synthesis to support viral protein glycosylation. Proc. Natl. Acad. Sci. 111, 18019–18024 (2014).

90. Wang, J. et al. Quantifying the RNA cap epitranscriptome reveals novel caps in cellular and viral RNA. Nucleic Acids Res. 47, e130–e130 (2019).

91. Mahajan, S., Choudhary, S., Kumar, P. & Tomar, S. Antiviral strategies targeting host factors and mechanisms obliging +ssRNA viral pathogens. Bioorg. Med. Chem. 46, 116356 (2021).

92. Ahmed, S. K. et al. Targeting Chikungunya Virus Replication by Benzoannulene Inhibitors. J. Med. Chem. 64, 4762–4786 (2021).

93. Yoon, J. et al. Design, Synthesis, and Anti-RNA Virus Activity of 6′-Fluorinated-Aristeromycin Analogues. J. Med. Chem. 62, 6346–6362 (2019).

94. Kumar, R. et al. S-adenosylmethionine-dependent methyltransferase inhibitor DZNep blocks transcription and translation of SARS-CoV-2 genome with a low tendency to select for drug-resistant viral variants. Antiviral Res. 197, 105232 (2022).

95. Chen, S. et al. Enhancer of zeste homolog 2 is a negative regulator of mitochondria-mediated innate immune responses. J. Immunol. 191, 2614–2623 (2013).

96. Tseng, C. K. H. et al. Synthesis of 3-deazaneplanocin A, a powerful inhibitor of S-adenosylhomocysteine hydrolase with potent and selective in vitro and in vivo antiviral activities. J. Med. Chem. 32, 1442–1446 (1989).

97. Moser, L. A. et al. A universal next-generation sequencing protocol to generate noninfectious barcoded cDNA libraries from high-containment RNA viruses. Msystems 1, e00039–15 (2016).

98. Franz, S. et al. Susceptibility of Chikungunya Virus to Inactivation by Heat and Commercially and World Health Organization-Recommended Biocides. J. Infect. Dis. 218, 1507–1510 (2018).

99. Lai, Y.-H., Franke, R., Pinkert, L., Overwin, H. & Brönstrup, M. Molecular Signatures of the Eagle Effect Induced by the Artificial Siderophore Conjugate LP-600 in E. coli. ACS Infect. Dis. 9, 567–581 (2023).

100. Su, D. et al. Quantitative analysis of ribonucleoside modifications in tRNA by HPLC-coupled mass spectrometry. Nat. Protoc. 9, 828–841 (2014).

101. Zheng, G. et al. Efficient and quantitative high-throughput tRNA sequencing. Nat. Methods 12, 835–837 (2015).

102. Livak, K. J. & Schmittgen, T. D. Analysis of relative gene expression data using real-time quantitative PCR and the 2− ΔΔCT method. methods 25, 402–408 (2001).

